# The PilT retraction ATPase promotes both extension and retraction of the MSHA type IVa pilus in *Vibrio cholerae*

**DOI:** 10.1101/2022.05.31.494186

**Authors:** Hannah Q. Hughes, Nicholas D. Christman, Triana N. Dalia, Courtney K. Ellison, Ankur B. Dalia

## Abstract

Diverse bacterial species use type IVa pili (T4aP) to interact with their environments. The dynamic extension and retraction of T4aP is critical for their function, but the mechanisms that regulate this dynamic activity remain poorly understood. T4aP are typically extended via the activity of a dedicated extension motor ATPase and retracted via the action of an antagonistic retraction motor ATPase called PilT. These motors are generally functionally independent, and loss of PilT commonly results in T4aP hyperpiliation due to undeterred pilus extension.

However, for the mannose-sensitive hemagglutinin (MSHA) T4aP of *Vibrio cholerae*, the loss of PilT results in a loss of surface piliation, which is unexpected based on our current understanding of T4aP dynamics. Here, we employ a combination of genetic and cell biological approaches to dissect the underlying mechanism. Our results demonstrate that PilT is necessary for MSHA pilus extension in addition to its well-established role in promoting MSHA pilus retraction. Through a suppressor screen, we also provide genetic evidence that the MshA major pilin impacts pilus extension. Together, these findings contribute to our understanding of the factors that regulate pilus extension and describe a previously uncharacterized function for the PilT motor ATPase.

**AUTHOR SUMMARY:** Many bacteria use filamentous appendages called type IVa pili to interact with their environment. These fibers dynamically extend and retract through the activity of ATPase motor proteins. In most pilus systems, deletion of the retraction motor results in uninterrupted pilus extension, leading to hyperpiliation. However, in the MSHA pilus system of *V. cholerae*, deletion of the retraction motor, *pilT*, results in a decrease in the number of surface pili. Here, we show that PilT is unexpectedly required for MSHA pilus extension in addition to its defined role in promoting pilus retraction. These results extend our understanding of the complex mechanisms underlying the dynamic activity of these broadly conserved filamentous appendages.

## INTRODUCTION

Type IVa pili (T4aP) are used by a wide range of bacteria to interact with their environment [1]. These fibers extend and retract from cell surfaces to accomplish a wide range of behaviors, including twitching motility [2, 3], the uptake of DNA during natural transformation [4, 5], and initial attachment of cells to a surface [6–8]. The frequency of pilus dynamic activity and the number of surface pili maintained at any given time directly impacts the ability of the cell to perform pilus-related functions. For example, the *Neisseria meningitidis* pilus system has been characterized to perform different functions based on the number of surface pili, ranging from DNA uptake, where a single pilus is sufficient, to interaction with host cells, which optimally requires five pili [9]. Elucidating the regulation of pilus dynamic activity and pilus number is therefore crucial to understanding pilus function.

T4aP are primarily composed of repeating subunits of the major pilin, which are extruded from the inner membrane and assembled into a filament through the activity of a dedicated extension ATPase. The process is reversed through the activity of a retraction ATPase, commonly called PilT, which deposits major pilins back into the inner membrane [10]. In these dynamic pilus systems, the activity of the extension and retraction motors plays an important role in regulating surface piliation. Current models propose that these motors compete to interact with the T4aP pilus machine, such that the motor that interacts with the platform dictates the direction of dynamic activity [11, 12]. If the extension motor is inhibited, pilus retraction dominates and pilus number decreases [13–15]. Conversely, if the retraction motor is disrupted, extension occurs undeterred, resulting in hyperpiliation [5, 16–19]. However, these trends are not universally observed, indicating that alternate mechanisms of pilus regulation exist and contribute to surface piliation.

Deletion of *pilT* does not result in hyperpiliation of the T4P in *Francisella tularensis* [20], *Clostridium perfringens* [21], or *V. cholerae* mannose-sensitive hemagglutinin (MSHA) pili [22–25]. Indeed, for each of these systems, deletion of *pilT* results in a decrease in piliation. This observation is counterintuitive based on our current understanding of T4aP and suggests that T4aP dynamic activity may be regulated by more than a simple competition between extension and retraction motors. In this study, we use the MSHA T4aP of *V. cholerae* as a model system to investigate PilT-dependent surface piliation because of the extensive genetic and cell biological tools that have recently been developed to study the dynamic activity of these pili [7, 22, 23, 25, 26].

## RESULTS

*Vibrio cholerae* expresses two distinct T4aP systems: MSHA pili and competence pili (also known as ChiRP pili [27]). MSHA pili facilitate biofilm formation by promoting the initial attachment of cells to a surface [8, 27–29], while competence pili promote DNA uptake for horizontal gene transfer by natural transformation [4, 5]. The dynamic activity of both T4aP systems can be studied in live cells via the use of a major pilin cysteine knock-in mutation (*mshA^T70C^* for MSHA pili; *pilA^S67C^* for competence pili) and subsequent labeling with thiol-reactive fluorescent maleimide dyes [4, 7, 26]. Competence pili are highly dynamic and pilus retraction generally occurs immediately following extension. In contrast, MSHA pili are maintained in an extended state and retract in response to distinct stimuli [23]. One stimulus that induces MSHA pilus retraction is the imaging condition used to visualize fluorescently labeled pili. Although the molecular mechanism underlying this response remains unclear, it provides a simple approach to assess MSHA pilus retraction. While these T4aP have distinct extension motor ATPases (MshE for MSHA pili; PilB for competence pili), they both rely on the same ATPase motor, PilT, for efficient retraction (**Fig. 1A-B**), which is consistent with a number of previous studies [4, 22, 23].

**Fig. 1.**
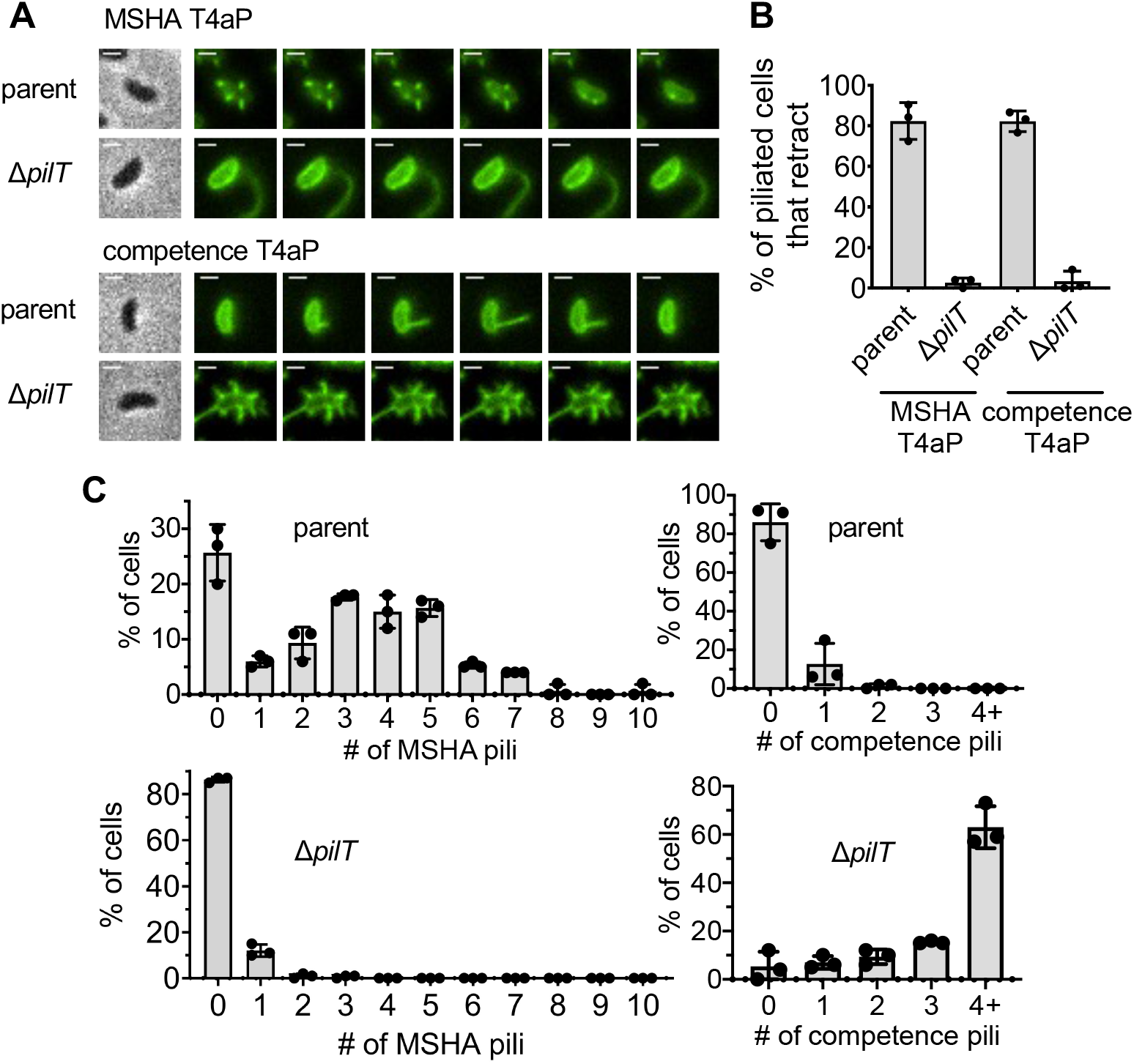
PilT has a distinct impact on *V. cholerae* MSHA and competence T4aP surface piliation. (**A**) Representative montage of piliated AF488-mal labeled cells for the pilus system indicated. Strains contain either *mshA^T70C^* (to label MSHA pili) or *pilA^S67C^* (to label competence pili). Phase images (left) are included from the first frame of the montage and show the cell boundary. Fluorescent images show AF488-mal labeled pili and there are 6-s intervals between frames. Scale bar = 1 μm. (**B**) Quantification of MSHA or competence T4aP retraction. Graph displays the percentage of piliated cells in each replicate that exhibited any pilus retraction during a three-minute timelapse. *n* ≥ 25 piliated cells analyzed for each of the three biological replicates. (**C**) Quantification of MSHA and competence pili in parent and Δ*pilT* cells as indicated. *n* = 300 cells analyzed from three independent biological replicates for all samples. All data are displayed as the mean ± SD.

Deletion of *pilT*, however, exhibits distinct effects on these two T4aP systems. Loss of *pilT* results in a marked increase in the number of surface competence pili, which is consistent with unregulated extension in this background as observed in many other T4aP systems [14, 30–33]. Conversely, the loss of PilT dramatically reduces the number of MSHA pili (**Fig. 1A, C**). In fact, in the Δ*pilT* mutant, most cells lack MSHA pili, and the rare cells that do make pili generally display a single pilus that is longer than those observed in the parent (**Fig. 1A, C**). This change in pilus number and length has been previously quantified and shown to differ significantly from the parent strain [23]. These phenotypes have also been qualitatively observed in multiple prior studies [4, 15, 22–25]. While the change in pilus length is visually striking, the major population-level change is the marked reduction in the frequency of piliated cells in the Δ*pilT* background (**Fig. 1C**).

One model that has been proposed for the observed reduction in MSHA surface piliation is that PilT is necessary to disrupt pilus extension after it is initiated [25]. Thus, in Δ*pilT* cells, a single MSHA T4aP machine undergoes uncontrolled “runaway” extension that exhausts the MshA major pilin pool of the cell. This would account for the markedly longer pili observed in the piliated Δ*pilT* cells (**Fig. 1A**). This model predicts that a similar proportion of cells should be piliated in both the parent and Δ*pilT* strains, since initiation of extension would not be affected. However, as mentioned above, the Δ*pilT* strain shows a striking reduction in the percentage of piliated cells (**Fig. 1C**). One possible explanation for this discrepancy could be that the long MSHA pili in the Δ*pilT* cells are sheared off of cells. If runaway extension was followed by shearing and loss of surface pili, we would expect Δ*pilT* cells to exhibit a distinct reduction in the level of cell-associated MshA major pilin. However, Western blot analysis indicates that cell-associated MshA major pilin is similar in the parent and Δ*pilT* strains (**Fig. 2A**). In the T4aP systems of *Pseudomonas aeruginosa* and *Myxococcus xanthus*, major pilin expression is increased in Δ*pilT* cells due to feedback regulation by the PilS-PilR two-component system [34, 35]. *V. cholerae* lacks homologs of the PilS-PilR system, but it remains possible that Δ*pilT* cells also compensate for the loss of sheared pili by increasing expression of MshA major pilin. If so, the total amount of major pilins present in the cells + supernatant (*i.e.*, unassembled + assembled, shed pili) should be markedly increased in the Δ*pilT* strain compared to the parent. However, we found that MshA protein levels were similar in the parent and Δ*pilT* samples even when the supernatants were included (**Fig. 2A**). These results are not consistent with runaway extension of MSHA pili in Δ*pilT* cells. We next assessed whether Δ*pilT* cells were instead experiencing a biogenesis defect. If so, we would expect them to retain parental levels of intracellular major pilin. The Alexa Fluor 488-maleimide dye (AF488-mal) used to fluorescently label cysteine modified MshA major pilins (MshA^T70C^) can pass through the outer membrane of *V. cholerae* and label pilins retained in the inner membrane [26]. Thus, the cell body fluorescence of labeled cells can serve as a proxy for the concentration of unassembled major pilins. To ensure that comparisons in this assay were not obscured by the fact that some strains have extended pili while others do not, we first labeled cells with AF488-mal, induced pilus retraction (see Methods for details), and then analyzed the cell-associated fluorescence of unpiliated cells as a proxy for the concentration of MshA major pilin they possess in their inner membrane. We compared the cell-associated fluorescence of the parent strain, a Δ*pilT* strain, a Δ*mshE* strain, and a strain lacking the *mshA^T70C^* mutation. The Δ*mshE* mutant lacks the canonical extension ATPase and correspondingly cannot assemble pili (no surface pili are ever observed in this mutant background). It was therefore included as a control to demonstrate the cell body fluorescence of a strain that retains all of its MshA major pilins in the inner membrane. The strain lacking the *mshA^T70C^* mutation (*i.e.*, unlabelable) was included to indicate the background fluorescence observed in the absence of any labelable major pilins. We found that the cell body fluorescence of unpiliated AF488-mal labeled Δ*pilT* cells was more similar to the parent and Δ*mshE* strains than to the unlabelable strain (**Fig. 2B**). This suggests that unpiliated Δ*pilT* cells retain a significant concentration of MshA in their inner membrane, a finding inconsistent with the runaway extension model.

**Fig. 2.**
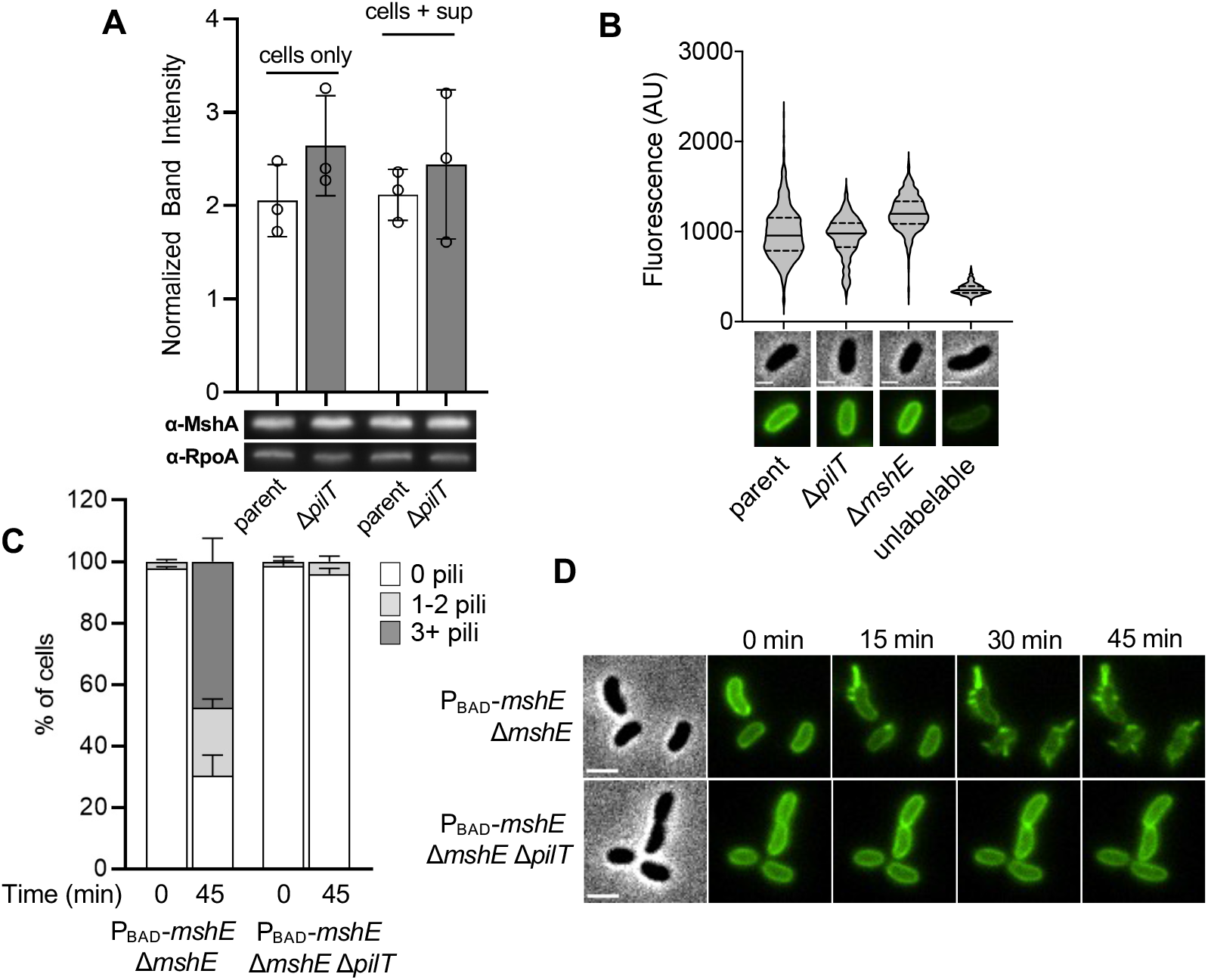
Reduced MSHA piliation of Δ*pilT* cells is likely due to a lack of pilus biogenesis rather than “runaway” extension. (**A**) Western blot analysis of MshA protein levels in parent and Δ*pilT* strains. Samples represent either cell-associated major pilin (cells only) or cell associated + pili shed in the supernatant (cells + sup). Band intensities are normalized to the RpoA loading control for quantification. Data are from three independent biological replicates. Data are displayed as the mean ± SD. (**B**) Quantification of cell body fluorescence in AF488-maleimide labeled cells as indicated. The parent, Δ*pilT*, and Δ*mshE* strains contain the *mshA^T70C^* mutation required for labeling the major pilin, while the unlabelable strain lacks this mutation. The latter is included here to denote the background cell body fluorescence observed in the absence of major pilins that can be labeled. The fluorescence of unpiliated cells was analyzed (as a measure of major pilin load in the inner membrane), and *n* ≥ 600 cells from three independent biological replicates for each strain. Data are presented as violin plots that demarcate the median (solid line) and quartile range (dotted lines). Representative images of cells used for quantification are included below. Phase images show cell boundaries and fluorescence images show AF488-mal labeled cells. Lookup tables are equivalent between fluorescence images. Scale bar = 1 μm. (**C**) Quantification of piliation in the indicated strains before and after induction with 0.2% arabinose. Cells were categorized as either having no pili (white bars), 1-2 pili (light gray bars), or at least 3 pili (dark gray bars). *n* = 300 cells analyzed from three independent biological replicates for all samples and data are displayed as the mean ± SD. (**D**) Representative montage of timelapse imaging of cells quantified in **C**. Scale bar = 2 µm.

Another way to test the runaway extension hypothesis is to directly observe pilus biogenesis and extension dynamics in parent and Δ*pilT* cells. Because the Δ*mshE* cells retained fluorescently labeled MshA in their inner membrane (**Fig. 2B**), we hypothesized that these AF488-mal labeled cells could be induced to extend their fluorescently labeled pilins into pilus filaments via ectopic induction of *mshE*. Furthermore, if *mshE* is induced in these cells under the microscope, this approach would circumvent any issues associated with pilus shearing, since pilus extension would be directly observed in single cells trapped under a gelzan pad. Therefore, if Δ*pilT* cells were undergoing runaway extension, we would expect most cells in the population to extend a single long pilus when *mshE* is induced. However, if the Δ*pilT* cells have an MSHA biogenesis defect, we would instead expect most cells in the population to remain unpiliated following *mshE* induction. To test this, we deleted the native copy of *mshE* and added an ectopic copy under the control of an arabinose-inducible promoter (P_BAD_-*mshE*). While parent cells (where *pilT* was intact) were able to extend pili after *mshE* was induced, Δ*pilT* cells showed a stark defect in pilus biogenesis (**Fig. 2C-D**), providing further evidence against the runaway extension model.

A lack of pilus biogenesis in the Δ*pilT* background could be due to an indirect downregulation of MshE extension motor activity. If so, we hypothesized that increased expression of *mshE* should recover surface piliation in the Δ*pilT* mutant. Overexpression of *mshE*, however, did not recover piliation in the Δ*pilT* background (**Fig. S1**). MshE activity is also allosterically regulated by cyclic-di-GMP (c-di-GMP), such that high levels of c-di-GMP stimulate MshE-dependent pilus extension [22, 36]. Thus, the lack of MSHA biogenesis in Δ*pilT* cells could be due to a decrease in cellular levels of c-di-GMP, diminishing MshE activity. To test this, we increased the intracellular concentration of c-di-GMP in Δ*pilT* cells by overexpressing a previously characterized diguanylate cyclase, DcpA [37]. Elevated c-di-GMP also induces expression of the *vps* and *rbm* loci [38], which encode *Vibrio* polysaccharide and biofilm matrix proteins, respectively. To avoid cell clumping due to the expression of these biofilm factors and for ease of pilus quantification in these experiments, we deleted both the *vps* and *rbm* loci (ΔVC0917-VC0939; here called Δ*vps*) in the background where *dcpA* was overexpressed. We found that overexpression of *dcpA* was insufficient to restore piliation in the Δ*pilT* background (**Fig. S1**). To further test whether c-di-GMP was involved in regulating MSHA piliation in the Δ*pilT* background, we took advantage of a previously-characterized allele of *mshE* – *mshE^L10A/L54AL/58A^* (denoted here as *mshE\**) – that genetically mimics the c-di-GMP bound state of MshE and promotes MSHA extension independently of c-di-GMP concentration [23, 36]. Native expression and overexpression of *mshE\**, however, both failed to overcome the MSHA surface piliation defect of the Δ*pilT* background (**Fig. S1**). These results suggest that PilT does not promote MSHA biogenesis by indirectly affecting MshE abundance or interaction with c-di-GMP.

Because PilT did not indirectly alter pilus number through known mechanisms of extension regulation, we hypothesized that PilT actively participates in MSHA biogenesis. If this were true, we would expect Δ*pilT* cells to extend MSHA pili very rapidly following PilT induction. To test this, we generated strains where *pilT* expression could be tightly controlled via an arabinose- and theophylline-inducible ectopic expression construct [39] (P_BAD_-ribo-*pilT*), and we measured piliation in cells before and after induction of the ectopic *pilT* construct. While only minor changes in piliation were observed in a strain still expressing native *pilT*, induction of ectopic *pilT* in the Δ*pilT* background rapidly recovered piliation to parental levels (**Fig. 3A-B**). This increase was not observed in strains lacking the P_BAD_-ribo-*pilT* construct, indicating that this response was not due to a nonspecific effect of the arabinose and theophylline inducers (**Fig. 3A-B**). Importantly, this effect was extremely rapid (within minutes) and occurred in single cells prior to cell division, using existing, prelabeled pilins (**Fig. 3, S2**). It is therefore unlikely that PilT is indirectly promoting biogenesis through regulation of T4aP machine assembly, because the structural elements of the T4aP machine are thought to be inserted into the cell envelope through cell growth and division [40]. Indeed, a fluorescent fusion to MshJ, a component of the MSHA T4aP machine, demonstrated that the number of MSHA machines was similar in the parent and Δ*pilT* backgrounds (**Fig. 3C**). Because ectopic expression of *pilT* in these experiments stimulated MSHA biogenesis, these data also further confirm that Δ*pilT* cells are not simply unpiliated due to runaway extension. Together, these data suggest that in addition to its defined role in promoting MSHA retraction, PilT also participates in the biogenesis of MSHA pili.

**Fig. 3.**
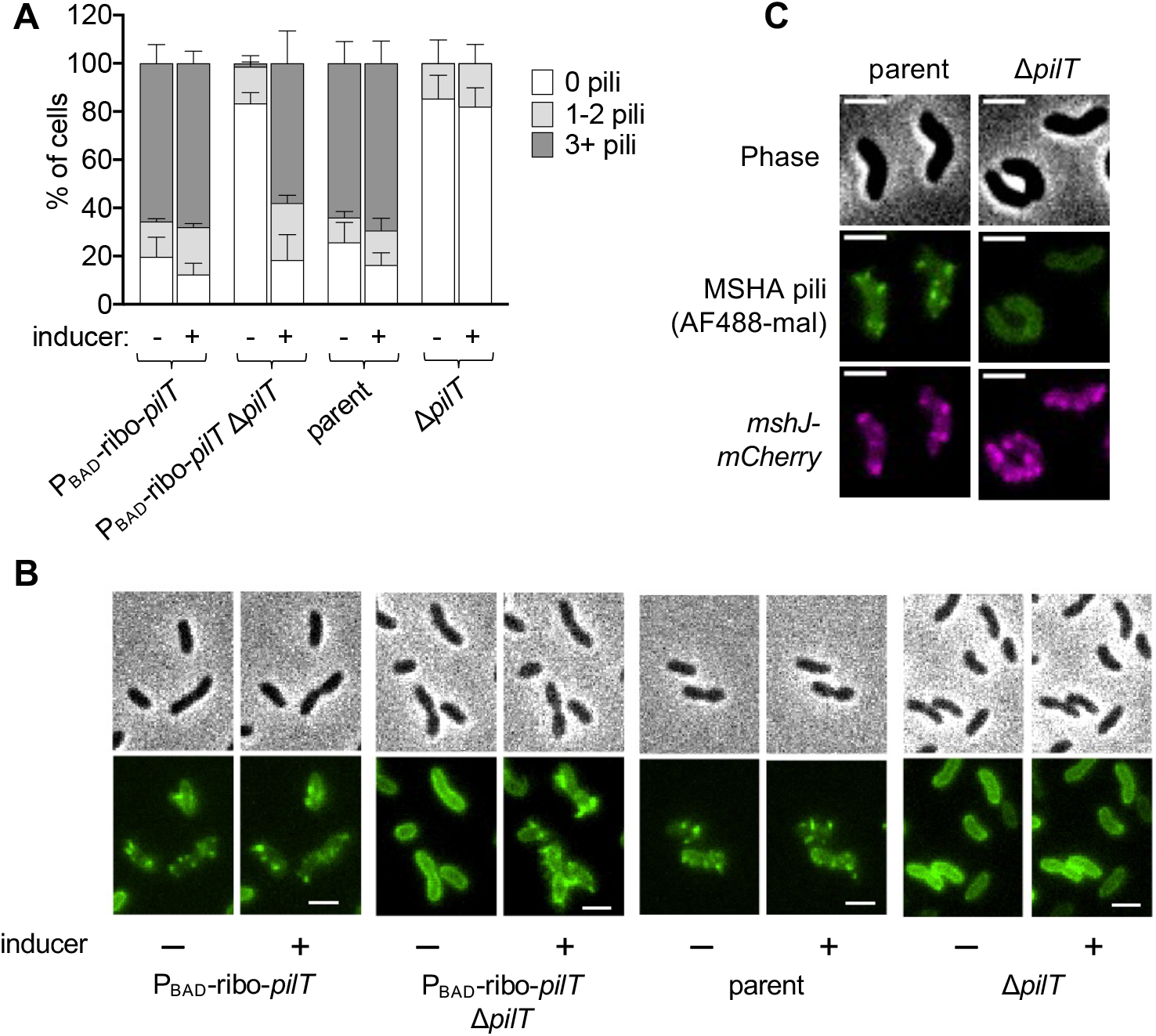
PilT promotes MSHA pilus extension. (**A**) Quantification of piliation in strains imaged before and after induction with 0.2% arabinose + 1.5 mM theophylline. Cells were categorized as either having no pili (white bars), 1-2 pili (light gray bars), or at least 3 pili (dark gray bars). *n* = 300 cells analyzed from three independent biological replicates for all samples and data are displayed as the mean ± SD. (**B**) Representative images of cells from strains in **A**. Phase images (top) show cell boundaries and fluorescence images (bottom) show AF488-mal labeled pili. Each sample is shown before (“-”) and after (“+”) induction. Scale bar = 2 μm. (**C**) Representative images of parent and Δ*pilT* strains containing a functional MshJ-mCherry fusion to label MSHA T4P machines. Phase images (top) show cell boundaries, green images (middle) show AF488-mal labeled pili, and red images (bottom) show MshJ-mCherry localization. Scale bar = 2 μm.

Some T4aP pili can undergo motor-independent retraction via the spontaneous depolymerization of the pilus filament [41]. Taking this into account, there are at least two distinct mechanisms by which PilT could support MSHA biogenesis. PilT can either (1) help promote MSHA pilus assembly, or (2) it can help maintain extended MSHA pili on the surface. For the latter model to be true, Δ*pilT* cells must be continually undergoing extension to generate short pili that cannot be resolved by our fluorescent imaging approach, and the rapid motor-independent retraction of these short filaments in the absence of PilT prevents the processive extension necessary to generate the surface pili visible in the parent strain. Under this model, once these pili are extended, PilT acts to prevent motor-independent retraction of these pili. If PilT is needed to maintain extended MSHA pili in this manner, we hypothesized that depleting PilT protein from piliated cells should result in the loss of surface pili due to the subsequent motor-independent retraction of those pili. The depletion of PilT was accomplished by adapting a previously described orthogonal degron system from *Mesoplasma florum* [42, 43]. The translational fusion of a degron tag (called pdt2) to the C-terminus of native PilT (*i.e.*, PilT-pdt2) allows for the specific and temporally regulated degradation of PilT by induction of the cognate *M. florum* Lon protease (mf-Lon). Strains with *pilT*-pdt2 exhibited parental levels of piliation without mf-Lon expression, while still exhibiting imaging-induced retraction, indicating that this fusion protein is functional (**Fig. S3A-C**). Importantly, depletion of PilT in these experiments is performed in conditions where cells are not actively dividing, which allows us to assess the effect of PilT depletion on cells with pre-extended pili, after initial biogenesis has already taken place. Piliation in strains with untagged PilT did not change upon mf-Lon induction, indicating that this protease does not pleiotropically affect MSHA piliation. The PilT-pdt2 strain also showed no change in piliation upon induction of the mf-Lon protease (**Fig. S3A-B**), indicating that the loss of PilT does not result in the loss of surface pili via motor-independent retraction. To assess whether PilT-pdt2 was indeed degraded in these experiments, we measured PilT-dependent retraction following depletion. Using imaging-induced retraction, we found that all strains exhibited normal retraction except for the PilT-pdt2 strain with induced mf-Lon (**Fig. S3C**), which is consistent with the depletion of PilT in this sample. Together, these data indicate that MSHA pili do not exhibit spontaneous retraction in the absence of PilT. Thus, PilT is not required to maintain extended pili, but instead helps to promote MSHA pilus biogenesis.

Because PilT is necessary for both MSHA extension and retraction, we next wanted to investigate whether both activities relied on its ATPase activity. In addition to PilT, *V. cholerae* also expresses PilU, a PilT-dependent accessory retraction motor ATPase. Loss of PilU alone (Δ*pilU*) did not impact MSHA pilus biogenesis, and the loss of PilU did not further exacerbate the piliation defect of the Δ*pilT* mutant (Δ*pilTU*) (**Fig. 4A-B, Fig. S4**). PilU is incapable of mediating retraction in the absence of PilT, but it can promote retraction in the presence of an ATPase defective allele of PilT [15, 25]. To test if PilTU-dependent MSHA retraction and extension were linked, we also assessed piliation in the ATPase defective Walker A mutants of PilT (*pilT^K136A^*) and PilU (*pilU^K134A^*). Although the *pilT^K136A^* allele is ATPase-defective, retraction can still occur in this background through the ATPase-activity of PilU (**Fig. 4C**). While the retraction-capable *pilT^K136A^* and *pilU^K134A^* mutants exhibited parental levels of MSHA piliation, the retraction-deficient *pilT^K136A^* Δ*pilU* and *pilT^K136A^ pilU^K134A^* mutants exhibited a loss of MSHA piliation (**Fig. 4A-C**). These data suggest that an ATPase active retraction motor – whether that is (1) PilT alone, or (2) a PilTU complex with functional PilU when PilT ATPase activity is compromised – is sufficient to promote MSHA pilus assembly.

**Fig. 4.**
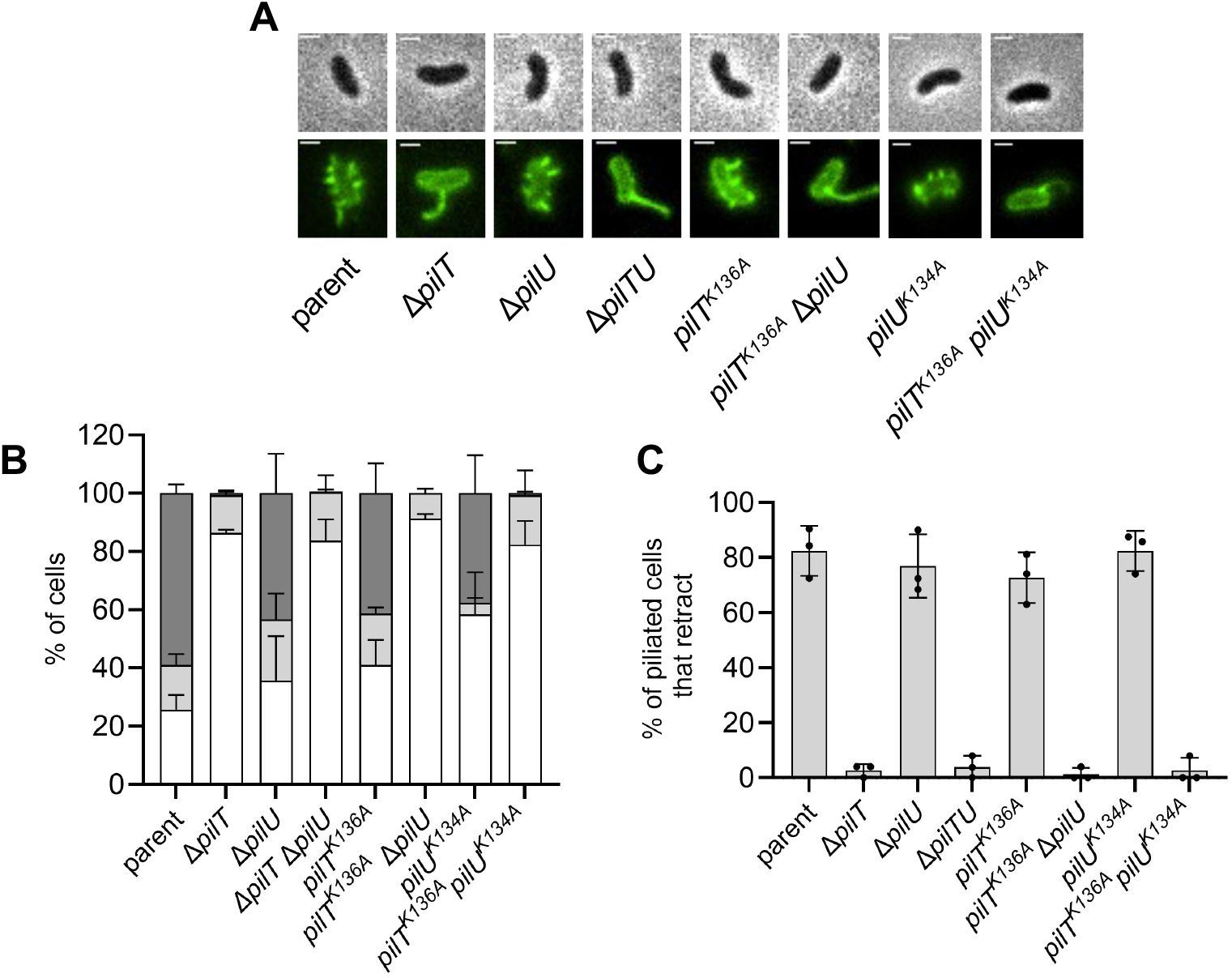
Retraction motor ATPase activity is required for MSHA surface piliation. (**A**) Representative images of piliated cells containing the indicated mutations to their retraction motor ATPases. Phase images (top) show cell boundaries and fluorescence images (bottom) show AF488-mal labeled pili. Scale bar = 1 μm. (**B**) Quantification of piliation in strains from **A**. Cells were categorized as either having no pili (white bars), 1-2 pili (light gray bars), or at least 3 pili (dark gray bars). *n* = 300 cells analyzed from three independent biological replicates for all samples. Data for the parent and Δ*pilT* strains are identical to that presented in **Fig. 1C** and are included again here for ease of comparison. (**C**) Quantification of retraction in strains from **A**. Graphs display the percentage of cells in each replicate that exhibited any pilus retraction during a three-minute timelapse. *n* ≥ 25 piliated cells analyzed for each of the three biological replicates. All data are displayed as the mean ± SD. Parent and Δ*pilT* data from **Fig. 1C** are included for comparison.

Our data thus far are consistent with a model wherein PilT promotes MSHA extension. We hypothesized that if PilT contributes to MSHA extension, it may directly interact with the MshE extension motor in order to carry out this function. If this interaction is specific to the role of PilT in extension of MSHA pili, we would predict no such interaction between PilT and the competence pilus extension motor, PilB, since PilT is not required for biogenesis of competence T4aP (**Fig. 1**). Because our data suggest that PilU can promote MSHA extension in the *pilT^K136A^* background, we also wanted to assess whether this accessory retraction ATPase exhibited a specific interaction with the MSHA extension motor. To assess interactions between these motors, we carried out bacterial adenylate cyclase two-hybrid (BACTH) analysis of the retraction motors (PilT and PilU) with the MSHA extension motor MshE and the competence pilus extension motor PilB. Interactions of the retraction motors with the MSHA platform protein (MshG) were also included as a positive control. The BACTH analysis showed that both PilT and PilU interacted with the MshG platform protein as expected (**Fig. S5**). Additionally, we found that the retraction motors (PilT and PilU) do interact with MshE (**Fig. S5**). However, these assays also showed interactions between the retraction motors and PilB (**Fig. S5**). Because the interaction between extension and retraction motors is not specific to the MSHA system, the physiological relevance for these interactions is unclear.

Thus far, our data suggest that PilT is required for MSHA pilus assembly. However, its precise role in the process is not yet established. One possibility is that PilT actually drives processive pilus extension via its ATPase activity. It has been shown in a number of T4P systems that the ATPase activity of the motor that drives dynamic activity correlates with the speed of extension/retraction [15, 41, 43, 44]. Thus, if PilT’s ATPase activity drives processive extension, we hypothesized that a mutation that slows its ATPase activity should correspondingly slow down extension speed. Previous work has defined mutant alleles of T4P motor ATPases that reduce ATPase activity ~1.5-2-fold [15, 41, 43, 45]. Therefore, we generated mutants of both *pilT* (*pilT^L201C^*; aka *pilT*^slow^) and *mshE* (*mshE^L390C^*; aka *mshE*^slow^) to slow their ATPase activity, and we tested the impact of these mutations on MSHA extension speed. MSHA pilus extension, however, is rarely observed in parent cells because the imaging conditions used to visualize pili induce MSHA retraction, as previously discussed. To counteract this limitation, we serendipitously found that some strains of *V. cholerae* are less susceptible to imaging-induced MSHA retraction. Namely, while the E7946 strain largely employed in this study is highly susceptible to imaging-induced retraction, the A1552 strain is much less sensitive. The A1552 strain exhibits the same loss of surface piliation when *pilT* is deleted, suggesting that the activity of PilT that promotes MSHA assembly is conserved across these *V. cholerae* strains [22]. Therefore, we employed the A1552 background to study pilus extension in these experiments. To facilitate direct observation of pilus extension, we generated strains where *mshE* could be induced while cells were imaged via timelapse microscopy (Δ*mshE* native + ectopic *mshE* expression construct). These strains also lacked native *pilU*, thereby eliminating any confounding effects this accessory motor ATPase might have on extension speed. When we assessed pilus extension speed in these strains, we found that while *mshE*^slow^ did reduce extension speed, *pilT*^slow^ had no impact on extension speed (**Fig. 5A**). Furthermore, cells that harbored both *mshE*^slow^ *pilT*^slow^ exhibited an extension speed that was indistinguishable from *mshE*^slow^ (**Fig. 5A**). These data suggest that PilT does not drive processive extension of MSHA pili. Instead, the canonical extension ATPase in this system, MshE, is responsible for this activity.

**Fig. 5.**
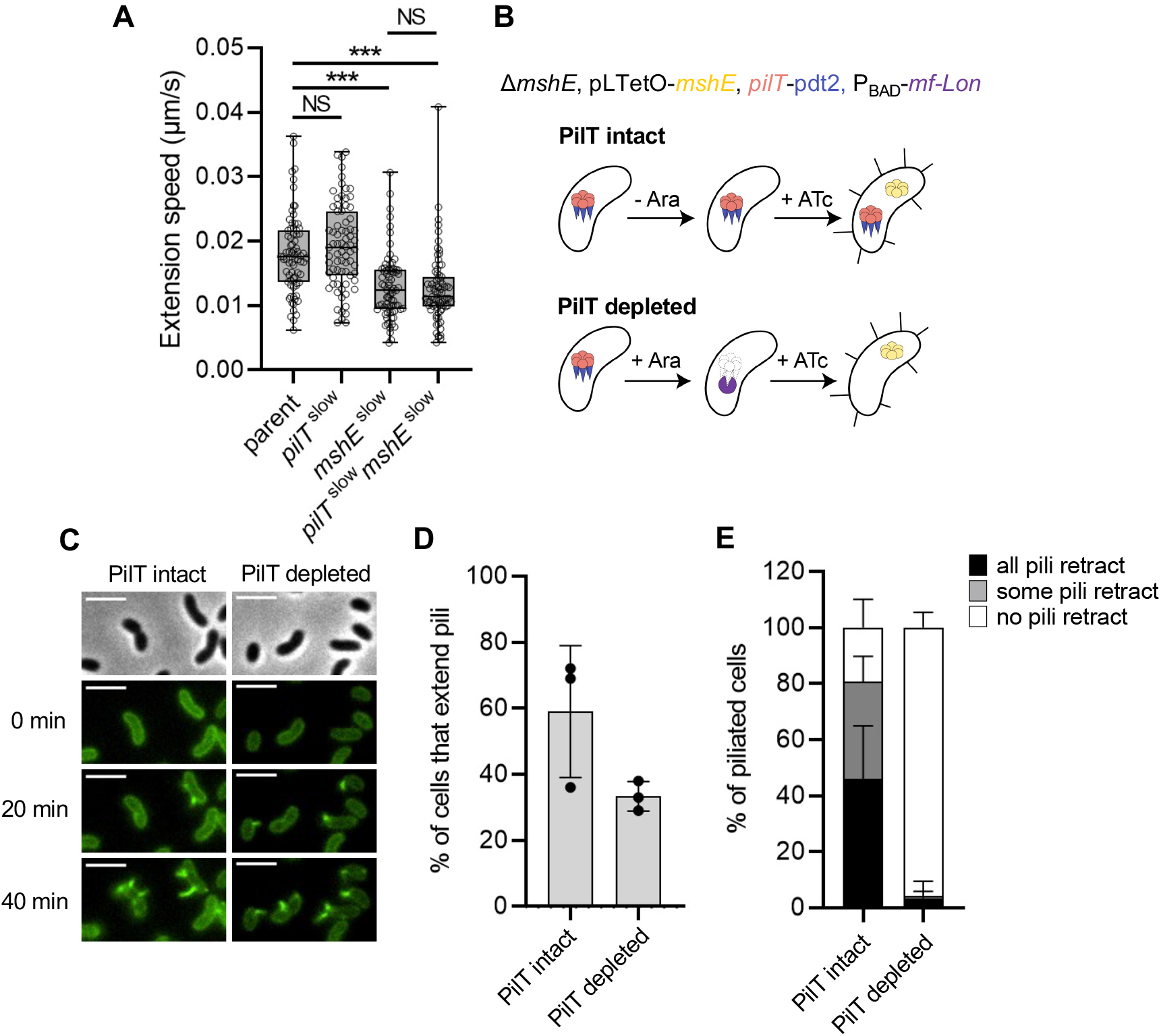
PilT and MshE promote distinct activities during MSHA extension. (**A**) The speed of MSHA T4P extension was determined in strains containing native motor ATPases (parent), or mutants that slow the ATPase activity of PilT (*pilT* ^slow^) and/or MshE (*mshE* ^slow^) as indicated. Because PilU is not required for MSHA piliation, all strains contained a Δ*pilU* mutation to prevent any confounding effects this motor ATPase might have on extension speed. For all strains, *n* = 75 from three independent experiments. Each data point indicates the extension speed for a dynamic T4P event. Box plots represent the median and the upper and lower quartile, while the whiskers demarcate the range. Statistical comparisons were made by One way ANOVA with Tukey’s post-test. NS = not significant; *** = *p* < 0.0001. (**B**) Schematic of the experimental setup to test whether PilT and MshE activity can be temporally separated in cells. In the absence of arabinose (Ara) induction (top), PilT-pd2 is left intact during *mshE* induction via anhydrotetracycline (ATc)(“PilT intact”). When cells are incubated with arabinose (bottom), PilT-pdt2 is degraded by Mf-Lon prior to the expression of *mshE* (“PilT depleted”). (**C**) Representative montage of cells subjected to the experiment described in **B**. Cells were imaged at 0, 20, and 40 minutes after induction of *mshE* expression. Scale bar = 3 μm. (**D**) Quantification of extension in the samples shown in **C**. Graph displays the percentage of cells in each replicate that exhibited pilus extension during a 40-minute window. *n* ≥ 100 cells analyzed for each of the three biological replicates and data are shown as the mean ± SD. (**E**) Quantification of imaging-induced retraction in samples shown in **C**. Graph displays the percentage of cells in each replicate that exhibited any pilus retraction during a three-minute timelapse. *n* ≥ 25 piliated cells analyzed for each of the three biological replicates and data are shown as the mean ± SD.

Our data above suggest that PilT does not actively drive processive extension. This result is also consistent with the observation that Δ*pilT* cells occasionally generate a single long pilus. This indicates that processive extension can still occur, albeit infrequently, even when PilT is absent. Based on these observations, we hypothesized that PilT may instead help initiate pilus extension. In canonical T4aP, pilus extension is initiated (or primed) by a complex of minor pilins - proteins that structurally resemble the major pilin but are expressed at significantly lower levels. One possibility is that the MSHA T4aP system minor pilins are not naturally capable of initiating pilus extension without the aid of PilT, which may help these minor pilins assemble and/or mature into a complex that is capable of initiating pilus assembly. If this were the case, PilT and MshE would be acting at distinct steps during MSHA pilus assembly and we would hypothesize that they would not need to be in the cell at the same time to promote extension. To test this, we used a strain where PilT-pdt2 could be degraded via ectopic induction of mf-Lon (P_BAD_-mf-Lon) and where *mshE* expression was under the control of a tightly inducible promoter (pLTetO-*mshE*). In these experiments, we grew cells without any inducers (so *mshE* was not expressed and pili were not extended) and then incubated them with AF488-mal to fluorescently label the MshA major pilin in their inner membrane. Next, these labeled cells were incubated either with or without arabinose to induce mf-Lon, which, if expressed, would deplete PilT-pdt2 from cells (**Fig. 5B**). Importantly, the depletion of PilT in these experiments is performed in conditions where cells are not actively dividing. Thus, the effect that PilT exerts on cells prior to depletion could theoretically be maintained (*i.e.*, if PilT facilitated maturation and/or assembly of the minor pilins into a complex) (**Fig. 5B**). Following this incubation, cells were placed under gelzan pads containing anhydrotetracycline (ATc), which induces *mshE* expression, and pilus extension was assessed via microscopy. If PilT and MshE activity can be temporally separated, we would expect cells to still be able to extend pili even when PilT was depleted. When we performed this experiment, we found that many cells could still extend pili, even when PilT was depleted (**Fig. 5C-D**). The percent of cells that extend pili is slightly lower when PilT is depleted (**Fig. 5C**). However, the amount of extension observed is still much greater than when cells lacked PilT altogether (**Fig. 2C**). The slightly reduced extension observed when PilT is depleted could be attributed to a short half-life for mature initiation complexes and/or due to the production of additional minor pilins while PilT was being depleted (and therefore could not be “matured” / assembled into initiation complexes). Importantly, we confirmed that piliated cells could not undergo imaging-induced retraction under the conditions where mf-Lon was induced, which phenotypically confirms that PilT-pdt2 is depleted in these experiments (**Fig. 5E**). All together, these results suggest that PilT and MshE activity can be temporally separated in cells. This is consistent with a model in which PilT facilitates initiation of pilus assembly, while MshE promotes processive extension. Furthermore, because the activity of PilT and MshE can be temporally separated, these data also reaffirm that PilT is not indirectly affecting MshE expression and/or activity.

To better understand factors regulating MSHA pilus extension, we also took an unbiased genetic approach. The presence of multiple surface pili is required for optimal MSHA-dependent attachment to surfaces [46]. Therefore, we took advantage of the poor attachment of Δ*pilT* cells to the walls of plastic culture tubes to select for suppressor mutants with improved binding. We hypothesized that this would occur through the restoration of MSHA surface piliation. Through this genetic selection, we isolated 14 spontaneous suppressor mutants that exhibited parental levels of binding and restored MSHA piliation. Whole genome resequencing revealed that all of the suppressor mutants had point mutations in the *mshA* major pilin, with 6 distinct *mshA* suppressor alleles identified (**Fig. S6A**). These suppressor mutations predominantly clustered around the predicted kink within the N-terminal α-helix (**Fig. 6A**). We reconstructed each of these mutations in clean genetic backgrounds, avoiding the use of rare codons, and saw that all of these *mshA* alleles increased MSHA piliation in the Δ*pilT* background (**Fig. 6B-C**). Correspondingly, these mutations also restored surface attachment to parental levels (**Fig. S6B**). Western blot analysis indicated that these alleles did not increase piliation by increasing the expression of the major pilin (**Fig. 6B**). To confirm that these *mshA* suppressor alleles relied on the canonical MSHA extension machinery, we also assessed piliation in a Δ*mshE* background. We found that none of the *mshA* suppressor alleles displayed surface pili in the Δ*mshE* background (*n* >1,000 cells analyzed per strain), which confirmed that extension of these *mshA* suppressor alleles was still dependent on the canonical MSHA extension machinery. Furthermore, all of these suppressor *mshA* mutants still exhibited PilT-dependent imaging-induced retraction, just like the *mshA*^parent^ (**Fig. 6D**). Thus, these suppressor mutations specifically enhanced MSHA surface piliation in the Δ*pilT* background without markedly altering MSHA piliation or retraction in the presence of *pilT*.

**Fig. 6.**
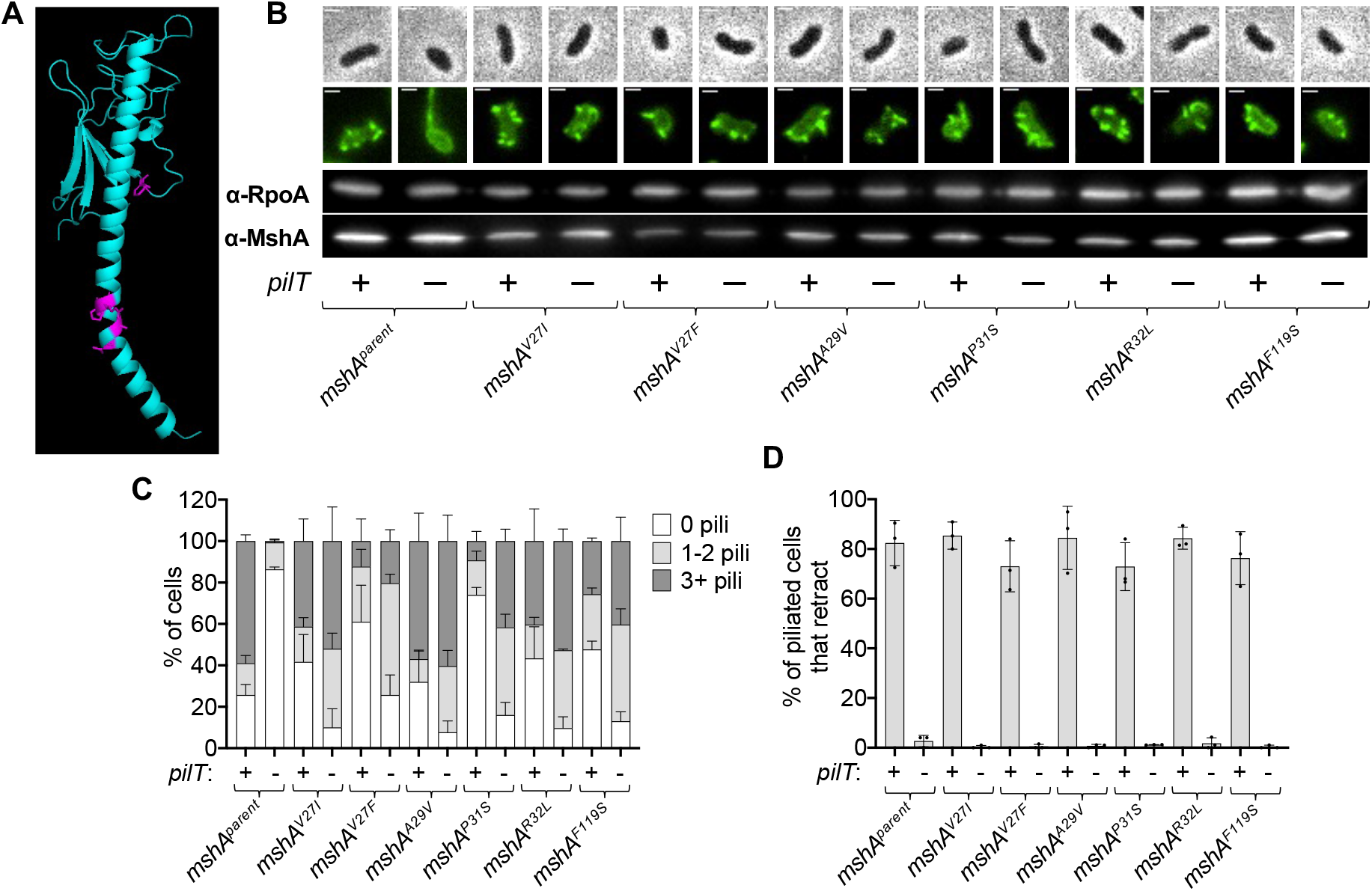
Point mutations in the major pilin *mshA* recover the piliation defect of a Δ*pilT* mutant. (**A**) Predicted structure of MshA, generated using AlphaFold [62, 63]. Residues mutated in the suppressor selection are colored magenta. (**B**) Representative images of piliated cells and Western blotting of MshA in suppressor mutants. RpoA is included as a loading control. Phase images (top) show cell boundaries and fluorescence images (below) show AF488-mal labeled pili. Scale bar = 1 μm. In all panels, strains that retain native *pilT* are denoted “+” and strains with Δ*pilT* mutations are denoted “-”. (**C**) Quantification of piliation in strains from **B**. Cells were categorized as either having no pili (white bars), 1-2 pili (light gray bars), or at least 3 pili (dark gray bars). *n* = 300 cells analyzed from three independent biological replicates for all samples. (**D**D) Quantification of retraction in strains from **B**. Data display the percentage of cells in each replicate that exhibited any pilus retraction during a three-minute timelapse. *n* ≥ 25 piliated cells analyzed for each of the three biological replicates. Data in **C**D and **D** are shown as the mean ± SD. Parent and Δ*pilT* data from **Fig. 1C** are included for comparison.

Since our previous data suggest that PilT helps initiate MSHA pilus extension, we hypothesized that the *mshA* suppressor mutants bypass the need for PilT by independently promoting pilus extension. As mentioned above, in canonical T4aP, minor pilins form an initiation complex that primes pilus assembly [47–49]. However, the MSHA minor pilins may not naturally be capable of initiating pilus extension without the aid of PilT. If so, the suppressor mutations in *mshA* could help enhance MSHA pilus extension in the absence of PilT by functioning similarly to a canonical T4aP minor pilin. As a result, we hypothesized that only low concentrations of the mutated MshA suppressor allele would be required to initiate pilus extension, since minor pilins are generally expressed at low levels. We chose *mshA^P31S^* as a representative suppressor because it was centrally located among the other suppressors, it was hit most frequently in the selection, and because we predicted that the disruption of the proline might generate the strongest impact on the structure of the protein. To test induction of low levels of *mshA^P31S^*, we generated constructs that ectopically expressed *mshA^P31S^* under the control of a tightly regulated P_tac_-riboswitch promoter, in a background that still expressed native *mshA* but lacked *pilT* (*i.e.*, *P_tac_*-ribo-*mshA^P31S^* Δ*pilT*). If MshA^P31S^ helps initiate pilus extension like a minor pilin, we hypothesized that it should be able to recover piliation even at low levels of induction. However, we found that full induction of the *mshA^P31S^* allele was required to increase pilus number (**Fig. S7A-B**). An equivalent construct inducing the wild-type *mshA* allele showed no change in pilus number, indicating that the observed effect on piliation requires ectopic expression of the *mshA^P31S^* suppressor allele. MshA expression from the P_tac_-riboswitch-*mshA* construct was assessed by Western blot analysis in a background that lacked native *mshA* (**Fig. S7C**), demonstrating that full induction resulted in native levels of MshA expression (**Fig. S7C**). Furthermore, we found that ectopic *mshA* induction in the presence of native *mshA* (*i.e.*, the scenario exhibited by the strains used in **Fig. S7B**) resulted in additive levels of MshA (**Fig. S7C**). These data suggest that the MshA suppressor alleles do not initiate pilus extension at low levels (akin to a canonical T4aP minor pilin). However, these alleles may still promote initiation of pilus assembly, albeit inefficiently.

If MshA^P31S^ does help initiate pilus extension in the absence of PilT, we hypothesized that the expression of this allele should promote assembly of wildtype MshA major pilins co-expressed in the same background. This cannot be distinguished in the experiments described above (**Fig. S7A-C**) because both the native *mshA* allele and the ectopically expressed *mshA* allele contain the T70C mutation that allows for fluorescent labeling. We therefore generated strains where only one of the two *mshA* alleles contained the T70C mutation, so that the assembly of each allele could be distinguished. If MshA^P31S^ can help initiate pilus assembly in the Δ*pilT* background, then we would expect ectopic expression of unlabeled MshA^P31S^ to promote the assembly of natively expressed, labelable MshA^T70C^ into fluorescent pilus filaments. Conversely, if MshA^P31S^ does not help initiate pilus assembly and instead assembles into distinct filaments, then no change in surface piliation should be observed when unlabeled MshA^P31S^ is induced. When we performed this experiment, we found that surface piliation was recovered following induction of MshA^P31S^ (native *mshA^T70C^* + ectopic *mshA^P31S^*) to a level that resembled the scenarios using labelable MshA^P31S^ (native *mshA^WT^* + ectopic *mshA^P31S,T70C^*; native *mshA^T70C^* + ectopic *mshA^P31S,T70C^*) (**Fig. 7**). Importantly, ectopic overexpression of the *mshA^parent^* alleles that lacked the P31S mutation (native *mshA^WT^* + ectopic *mshA^T70C^*; native *mshA^T70C^* + ectopic *mshA^WT^*) did not restore surface piliation (**Fig. 7**). Similar experiments using the *mshA^V27F^* and *mshA^R32L^* alleles also promoted assembly of labelable MshA^T70C^ into pilus filaments, suggesting this activity was not unique to the *mshA^P31S^* allele (**Fig. S8**). Together, these data indicate that these *mshA* suppressor alleles promote the assembly of *mshA^WT^* into pilus filaments, which is consistent with a model wherein the *mshA* suppressor alleles increase PilT-independent piliation by promoting initiation of pilus extension.

**Fig. 7.**
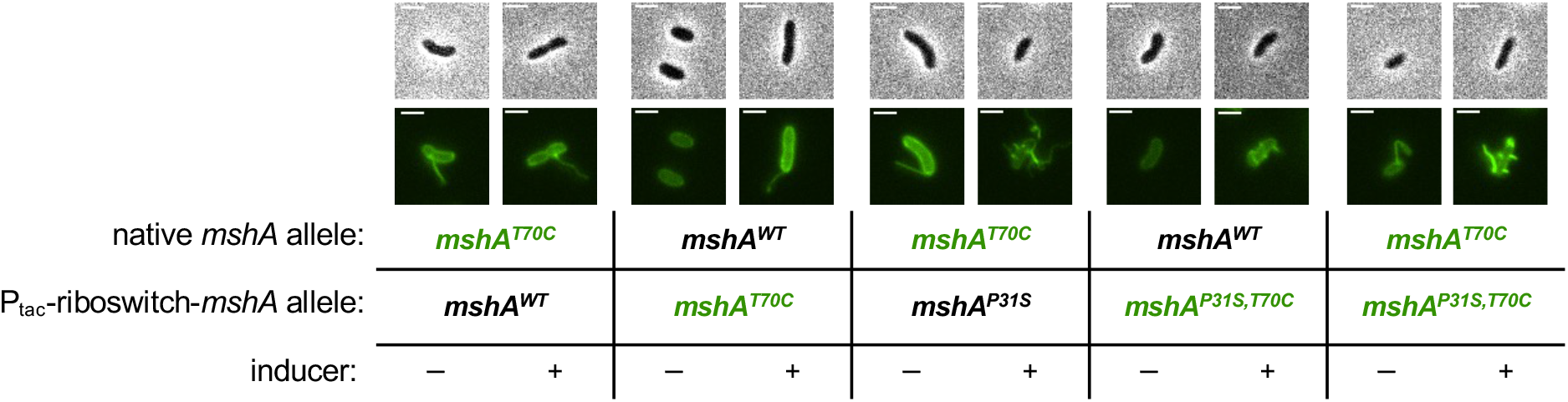
The MshA^P31S^ suppressor allele promotes assembly of wild-type MshA. Representative images of cells from Δ*pilT* strains expressing the indicated *mshA* allele at the native locus (native *mshA* allele) and ectopic locus (P_tac_-riboswitch-*mshA* allele). Cells were either grown with (”+”) or without (”-”) 100 µM IPTG + 1.5 mM theophylline to induce the ectopic P_tac_-riboswitch-*mshA* allele in the strain as indicated. Only the alleles containing the T70C mutation can be labeled with AF488-mal and are denoted in green text in the table below the images. Lookup tables are equivalent between fluorescent images to reflect the expression level of the labelable *mshA* allele. Scale bar = 2 μm.

## DISCUSSION

Here, we have explored the mechanistic basis for the regulation of MSHA piliation in *V. cholerae* and found that piliation is dependent on both PilT and the major pilin (**Fig. 8**). We demonstrate that PilT is not needed to disrupt runaway extension or maintain pili in their extended state. Instead, our data are consistent with a model in which PilT counterintuitively promotes MSHA pilus extension in addition to its established role in pilus retraction. The strongest evidence for this model comes from the observation that the tightly controlled ectopic expression of *pilT* in Δ*pilT* cells results in the rapid extension of MSHA pili and the observation that tightly controlled ectopic expression of *mshE* only recovers piliation when *pilT* is intact. Furthermore, we show that both PilT-dependent extension and retraction rely on the presence of a functional ATPase retraction motor, whether that is accomplished by PilT alone, or through the ATPase activity of PilU when PilT ATPase activity is inactivated.

**Fig. 8.**
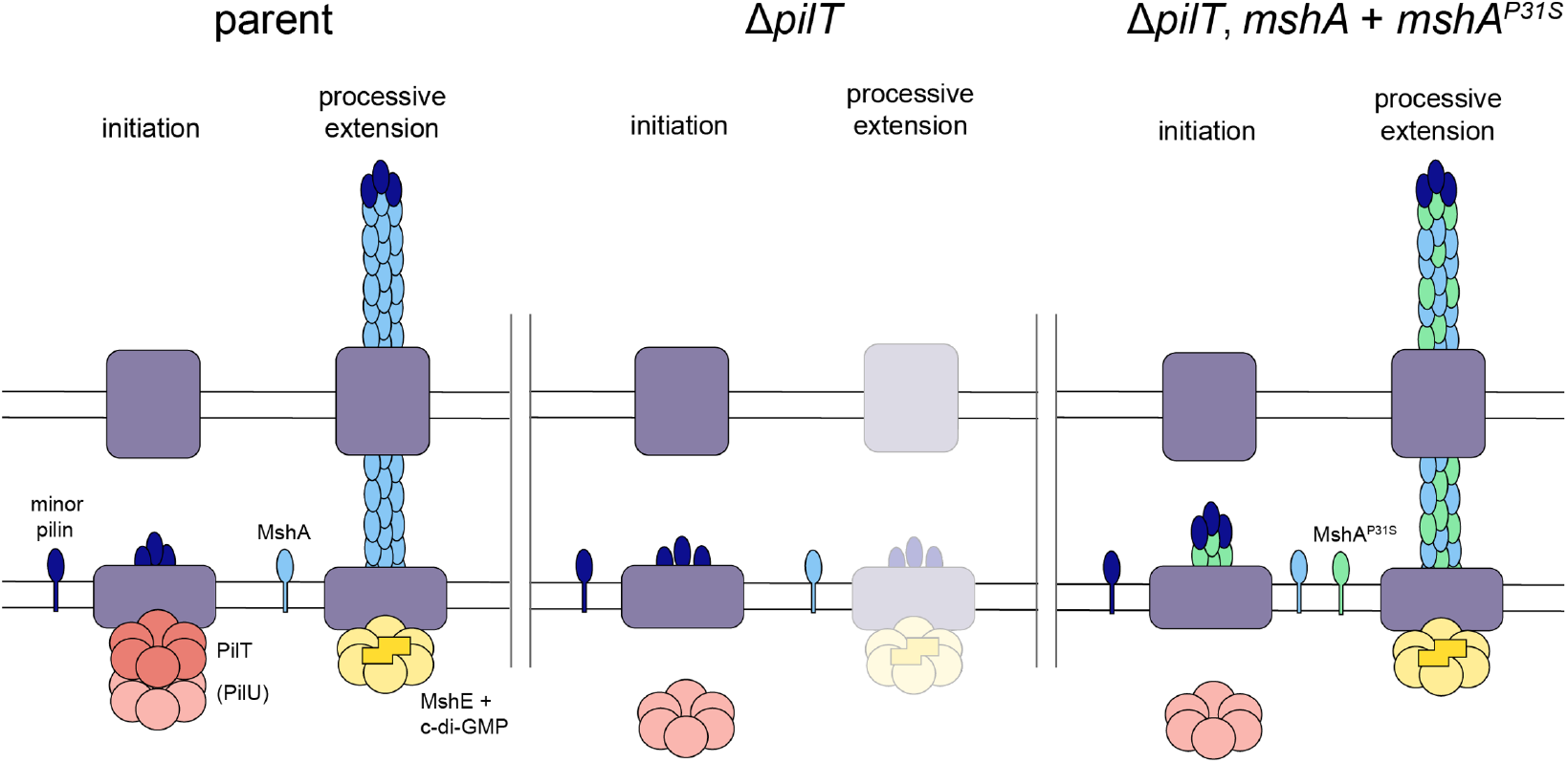
Model for the proposed mechanisms that promote MSHA extension and surface piliation. In wildtype cells (left), PilT promotes initiation of pilus extension, potentially by maturing the minor pilins into an initiation complex. Then, MshE can promote processive extension of the pilus in a c-di-GMP-dependent manner. The PilT-dependent accessory retraction ATPase motor PilU is dispensable for promoting pilus extension, but it can facilitate extension if PilT’s ATPase activity is inactivated. If PilT is absent (middle), initiation is inhibited, so processive extension is greatly diminished. Finally, mutations to the MshA major pilin (right), like MshA^P31S^, can promote initiation of MSHA extension even in the absence of PilT, suggesting that the structure of the major pilin is an important determinant of pilus extension.

We found that while PilT ATPase activity is required for MSHA assembly, PilT ATPase activity likely does not drive processive extension, and may not even need to be present during processive extension. These data are most consistent with a model wherein PilT is required for initiating pilus assembly (**Fig. 8**). In canonical T4aP, the minor pilins that initiate pilus assembly sit within the pilus machine prior to processive extension [12, 49]. So, one possibility is that PilT promotes retraction early on when only the minor pilins are sitting in the pilus machine to reorient the initiation complex (*i.e.*, the molecular equivalent of rotating a screw counterclockwise to ensure sure that the threads are seated properly). Alternatively, PilT could promote the interaction between minor pilins to generate a “mature” initiation complex. Studying the impact of minor pilins on MSHA pilus assembly and the potential role for PilT in facilitating their assembly will be the focus of future work.

Our results suggest that MSHA pilus biogenesis and dynamics may not simply be the result of direct competition between the extension and retraction motors in *V. cholerae*. Instead, we find that the combined activity of these motors is required for pilus extension. Recent work highlights that distinct motor ATPases can coordinate to promote pilus extension and/or retraction. As discussed above, PilU is an accessory retraction motor that works in conjunction with PilT to promote retraction of T4aP [15, 25]. In *Acinetobacter baylyi*, optimal T4aP extension depends on the combined activity of two distinct extension motors, TfpB and PilB, where TfpB initiates pilus extension and PilB promotes processive extension [14]. MSHA extension may be similarly regulated, through the cooperative activities of an extension and retraction motor (*i.e.*, MshE and PilT). Specifically, our data show that PilT is likely required for initiation of pilus extension, while MshE drives processive extension. PilT, however, is also clearly required for MSHA retraction. This bidirectional activity resembles *Caulobacter crescentus* CpaF, which is the sole motor ATPase expressed in this T4cP tad pilus system, and which drives both pilus extension and retraction [44]. Indeed, recent structural analysis of the transitions adopted by PilT suggest that this family of ATPase motors may be able to adopt distinct configurations that can switch the directionality of their activity (from retraction to extension) [50]. Because our data demonstrates that PilT ATPase activity does not directly drive extension speed, this distinguishes it from the bifunctional CpaF motor.

Our suppressor screen also implicated the structure of the major pilin in regulating MSHA biogenesis. Most of the suppressor mutations obtained were localized near a proline in the widely conserved N-terminal α-helix of T4aP major pilins (*i.e.*, the α1 subdomain) [51]. This proline introduces a kink into α1 [52], and is hypothesized to affect the packing of major pilin subunits within the filament. Indeed, mutations near the conserved proline in the major pilin of the *Myxococcus xanthus* T4aP have been shown to impact pilus assembly and/or retraction [53]. Because our data indicate that PilT promotes initiation of pilus extension, we hypothesize that these MshA suppressor alleles also enhance initiation of pilus extension to allow for pilus biogenesis in the absence of PilT. One possibility is that these suppressor alleles interact more efficiently with the minor pilin proteins so that pilus assembly does not depend on the “maturation” of the initiation complex carried out by PilT (**Fig. 8**). This is supported by the observation that the MshA suppressor alleles promote assembly of the MshA^parent^ major pilin into a pilus filament. Additionally, we have recently shown that the packing of major pilin subunits within the pilus filament plays an important role in regulating motor-independent retraction of T4aP [41]. This suggests that the major pilin and the structure of the pilus filament are factors that likely contribute to the regulation of both T4aP extension and retraction.

Ultimately, this study describes two unappreciated factors that influence pilus extension of the *V. cholerae* MSHA pili: the PilT retraction motor and the structure of the MSHA major pilin. Because PilT is also required for piliation in other T4aP systems [20, 21], it is likely that the mechanisms described here are not limited to the MSHA pili of *V. cholerae*. Also, recent phylogenetic analysis indicates that MSHA pilus systems form a discrete cluster from canonical T4aP [1]. In particular, MSHA systems contain pilus components of unknown function that are distinct from canonical T4aP (*i.e.*, MshM, MshQ, and MshF) [1]. It is possible that these pilus components contribute to PilT-dependent extension. Thus, characterizing the function of the pilus components that are unique to MSHA T4aP moving forward may yield further insight into the broad diversity of mechanisms that regulate pilus dynamic activity.

Ultimately, our findings describe a previously uncharacterized role for PilT. In addition to its canonical function as a retraction ATPase motor, we demonstrate that it is also required for the initiation of MSHA pilus assembly. These results expand our understanding of the factors that regulate the biogenesis and dynamic activity of these widely conserved bacterial surface appendages.

## METHODS

### Bacterial strains and culture conditions

*Vibrio cholerae* E7946 was used as the background for all strains employed in this study. Cultures were grown in LB broth (Miller) and plated on LB agar, and incubations were carried out at 30°C. Descriptions of all strains used in this study are found in **Supplemental Table 1**. Where appropriate, media were supplemented with kanamycin 50 μg/mL, trimethoprim 10 μg/mL, spectinomycin 200 μg/mL, chloramphenicol 1 μg/mL, carbenicillin 100 μg/mL, or zeocin 100 μg/mL.

### Construction of mutant strains

Splicing-by-overlap extension PCR was used to generate mutant constructs, as described previously [54]. Natural transformation and cotransformation were used to introduce mutant constructs into chitin-induced competent *V. cholerae* as described previously [55, 56]. Mutations were confirmed by PCR and/or sequencing. All ectopic expression constructs were derived from previously published strains [39]. The primers used to generate mutant constructs can be found in **Supplemental Table 2**.

### Microscopy and image analysis

All phase contrast and fluorescence images were obtained with a Nikon Ti-2 microscope with a Plan Apo 60X objective, GFP filter cube, Hamamatsu ORCAFlash 4.0 camera, and Nikon NIS Elements imaging software. Images were prepared with Fiji [57, 58].

Pili were visualized by Alexa Fluor 488-maleimide (AF488-mal) labelling. Cells were grown in LB to an OD_600_ of ~0.5 at 30°C rolling, with the exception of samples for imaging competence pili (which were grown in LB + 100 μM IPTG + 20 mM MgCl_2_ + 10 mM CaCl_2_ to an OD_600_ of ~2.5) or samples imaging *mshJ-mCherry* (which were grown in LB overnight). Cells were then washed and resuspended in instant ocean medium (7 g/L; Aquarium Systems) and incubated with 25 μg/mL AF488-mal for 10-15 min at room temperature. Cells were washed twice to remove excess dye and resuspended in instant ocean medium. After labelling, cells were loaded on a glass coverslip, covered with a 0.2% gelzan pad, and imaged. Lookup tables were chosen to best represent piliation for each image unless otherwise noted. Pilus number was assessed by randomly selecting 100 cells in a field of view in the phase channel for each replicate and then manually scoring piliation in each cell. The percent of piliated cells that exhibited retraction was assessed for each replicate by randomly selecting ~25 piliated cells and then observing retraction during a 3-minute timelapse (frames taken with 100 ms exposure every 3 sec). The retraction of any pili within a cell was used as a positive indicator of retraction. Three biological replicates were included for each dataset.

Extension speeds were obtained by first labelling cells as above. Cells were then loaded onto a glass coverslip and covered with an 0.2% gelzan pad containing 0.2% arabinose inducer to induce *mshE* expression. Timelapses (3 second intervals over 3 minutes) were taken ~15-30 minutes after addition of the inducer. Pilus length was measured in the final frame of the extension event, and extension speed was calculated by dividing that length by the total extension time.

To quantify cell body fluorescence, cells were labelled and imaged as described above, with the exception that cells were incubated with AF488-mal for 45 min. Pilus retraction (to ensure cells were unpiliated) was induced via imaging-induced retraction (*i.e.*, a 4-minute timelapse was performed where images were captured every 3 seconds). The final image of the timelapse was used for quantification of cell body fluorescence. Cell body detection and fluorescence quantification were carried out using MicrobeJ [59]. Data sets were manually curated to remove piliated cells, overlapping cells, and dead cells.

### Temporally regulated induction of pilT

Cells were first labelled, added to a coverslip, and covered with an 0.2% gelzan pad as described above. For **Fig. S2**, to observe how quickly cells could respond to inducer, cells were imaged under pads that either contained or lacked inducers (1.5 mM theophylline + 0.15% arabinose). An image was taken immediately upon exposure to the gelzan pad, and again after 15 min incubation at room temperature. For a more quantitative comparison of piliation following induction as shown in **Fig. 3**, cells were first loaded under a gelzan pad containing no inducer and allowed to recover for 5 min. A second 0.2% gelzan pad containing 3 mM theophylline + 0.4% arabinose was then layered on top of the first gelzan pad. The use of stacked gelzan pads in this experiment allowed for the slower diffusion of inducer molecules to cells (*i.e.*, via the diffusion of inducers through the gelzan pad in immediate contact with the cells). This ensured that any change in piliation was due to the effect of the inducer rather than recovery from conditions experienced during labeling and setup. Cells were imaged immediately (“-inducer”) and then again after 45 min, which we empirically determined was sufficient for the inducer to diffuse into the lower pad (“+ inducer”). Piliation states were quantified as above for each of the three biological replicates.

### Degradation of pilT

Strains were labelled as described above and then allowed to incubate for 2.5 hrs with either inducer (0.2% arabinose) or vector control (an equivalent volume of water) to induce mf-Lon. Cells were then loaded onto a coverslip, covered with an 0.2% gelzan pad, and imaged as described above. Piliation state was scored as described above. Retraction was scored over a 3 min timelapse as described above, but categorized into either “no pili retract”, “some pili retract”, or “all pili retract”. This was to reflect the more subtle effects seen during PilT depletion compared to when *pilT* is deleted.

For the experiment testing temporally distinct expression of MshE and PilT, cells were first labelled and PilT was degraded with 0.2% arabinose as described above. Cells were then loaded onto a coverslip and covered with a gelzan pad containing 50 ng/mL ATc to induce expression of *mshE*. Cells were imaged as above, after 0 min, 20 min, or 40 min in the presence of the inducer and scored for pilus extension. Pili were induced to retract through timelapse imaging and retraction was scored as described above.

### Western blotting

Cells were grown to an OD_600_ of ~0.5 at 30°C rolling. To assess cell associated MshA, cells were spun down (~1.5 × 10^9^ CFUs) and resuspended in an equivalent volume of instant ocean medium (*i.e.*, cells only). To assess the amount of MshA that was cell associated and shed, an aliquot of culture was used directly for Western blot analysis (*i.e.*, cells + sup). Samples were mixed 1:1 with 2 x SDS sample buffer [200 mM Tris pH 6.8, 25% glycerol, 1.8% SDS, 0.02% Bromophenol Blue, 5% β-mercaptoethanol] and then incubated at 95°C for 10 min, separated by SDS PAGE (15% gel), and then electrophoretically transferred to a PVDF membrane. This membrane was blocked with 5% milk and then incubated simultaneously with rabbit polyclonal α-MshA (1:1000, gift of J. Zhu) and mouse monoclonal α-RpoA (1:1000, Biolegend) (as a loading control) primary antibodies. Blots were washed and then incubated simultaneously with α-mouse and α-rabbit secondary antibodies conjugated to horseradish peroxidase enzyme. Blots were developed with Pierce™ ECL Western Blotting Substrate and then imaged with a ProteinSimple FluorChem R system.

### Bacterial adenylate cyclase two-hybrid assay

Genes of interest were amplified via PCR and cloned into the BACTH vectors pKT25 and pUT18C to produce N-terminal fusions between the target protein and the T25 and T18 fragments of adenylate cyclase, respectively. Miniprepped plasmids (Qiagen) were then co-transformed into *E. coli* BTH101 cells. As a positive control, T25-zip and T18-zip vectors were cotransformed into BTH101. As a negative control, the pKT25 and pUT18C empty vectors were cotransformed into BTH101. Transformations were plated onto LB + kanamycin (50 μg/mL) + carbenicillin (100 μg/mL) plates to select for transformants that received both plasmids. These transformants were then picked and grown overnight at 30°C in LB + kanamycin (50 μg/mL) + carbenicillin (100 μg/mL). Finally, 3 μL of the overnight cultures were spotted onto LB agar plates supplemented with 500 µM IPTG, kanamycin (50 μg/mL), carbenicillin (100 μg/mL), and X-gal (40 μg/mL). These plates were allowed to incubate statically at 30 °C for ~ 24 hours prior to imaging on an Epson Perfection V600 photo flatbed scanner.

### Genetic suppressor selection

20 independent cultures of Δ*pilT* Δ*vps* were started from ~100 CFUs each to generate independent lineages for this suppressor screen. Cells were grown in polystyrene culture tubes to an OD_600_ of ~1.5 at 30°C rolling. A small aliquot was removed from each culture tube to be used for quantitative plating on LB agar (total cells). Tubes were then dumped and rinsed 5 times with fresh LB to remove unbound cells. Bound cells were then removed from the walls of the culture tube by adding 3 mL of fresh LB medium and vortexing vigorously for 30 sec. The cell slurry was then centrifuged, concentrated, and subjected to quantitative plating on LB agar (bound cells). The binding frequency was calculated for each sample by dividing the CFU after rinsing (bound cells) by the CFU before rinsing (total cells). Cells retrieved after binding were used to inoculate the next round of selection. Iterative rounds of selection were continued for each lineage until the binding frequency was at least 100x higher than the parental Δ*pilT* Δ*vps* control (4-8 rounds selection). Genomic DNA from select clones was used to generate sequencing libraries by homopolymer tail mediated ligation PCR (HTML-PCR) exactly as previously described [60] and then sequenced on an Illumina Next-Seq instrument. The resulting sequencing data was mapped to the N16961 genome [61] using CLC Genomics WorkBench and the Basic Variant Analysis tool was used to identify SNVs.

### Binding assays

Cells were grown to an OD_600_ of ~2.5 at 30°C shaking and 4 mL of culture was loaded into a new polystyrene culture tube. Tubes were incubated on a roller drum at room temperature for 20 min, during which time cells were allowed to bind to the walls of the tube. After incubation, a small aliquot was removed from each culture tube to be used for quantitative plating on LB agar (total cells). Then, tubes were dumped and rinsed 5 times with fresh LB to remove unbound cells. Bound cells were then removed from the walls of the culture tube by adding 1 mL of fresh LB medium and vortexing vigorously for 30 sec. The cell slurry was then centrifuged, concentrated, and subjected to quantitative plating on LB agar (bound cells). The binding frequency was calculated for each sample by dividing the CFU after rinsing (bound cells) by the CFU before rinsing (total cells). Three biological replicates were carried out for each strain.

## ACKNOWLEDGEMENTS

We would like to thank Clay Fuqua, Julia van Kessel, and Tuli Mukhophadyay for helpful discussions. We would also like to thank Jay Zhu for providing α-MshA antibody. This work was supported by grant R35GM128674 (to ABD).

## AUTHOR CONTRIBUTIONS

H. Hughes, C. Ellison, and A. Dalia conceptualized the experiments. Investigation was carried out by H. Hughes, N. Christman, T. Dalia, and A. Dalia. H. Hughes and A. Dalia wrote the paper and incorporated feedback from all authors.

---

**Fig. S1.**
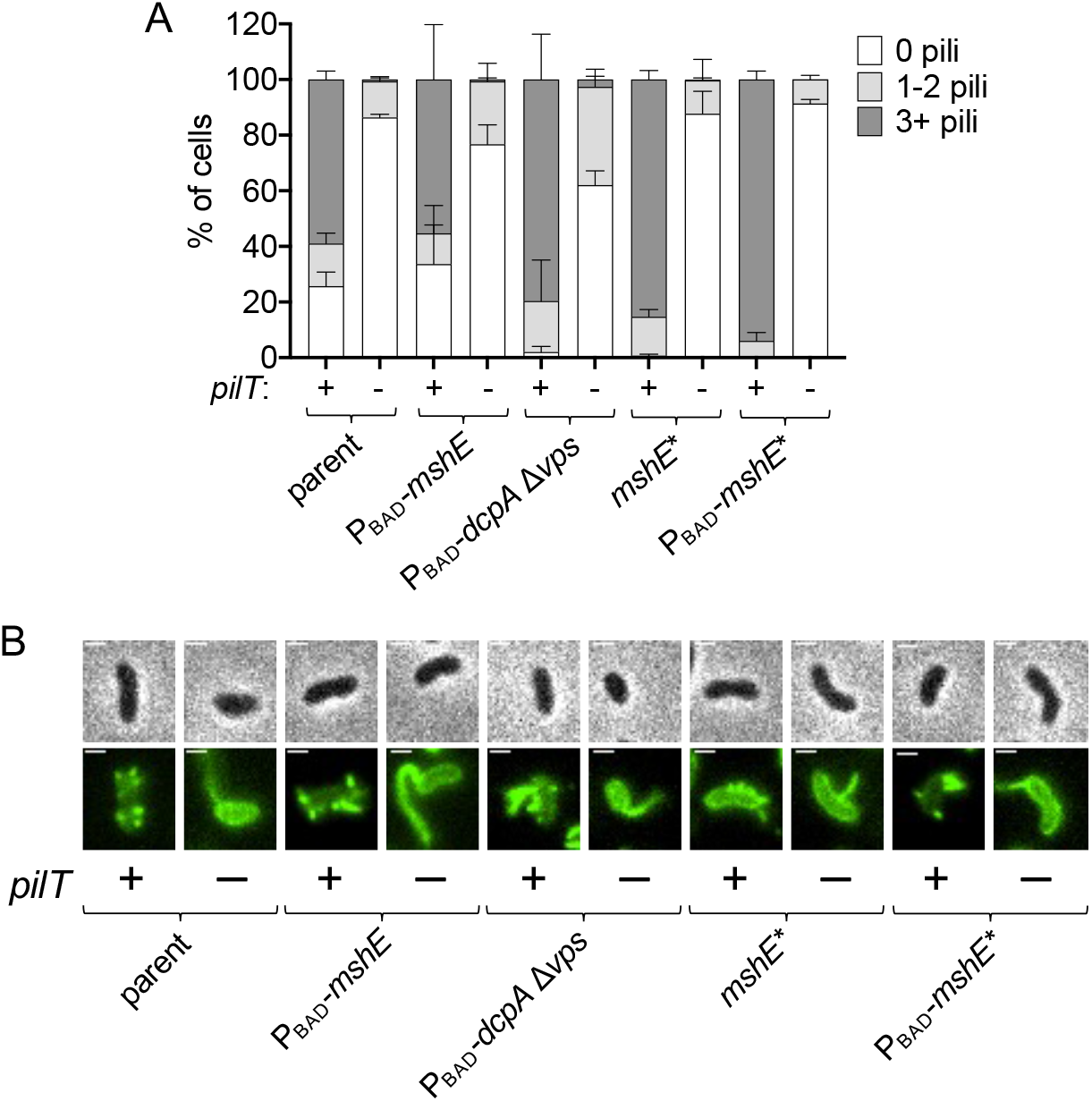
The piliation defect in the Δ*pilT* mutant cannot be recovered by increasing MshE expression or activity. (**A**) Quantification of piliation in strains with altered extension motor regulation. Strains that retain native *pilT* are denoted “+” and strains with Δ*pilT* mutations are denoted “-”. Cells were categorized as either having no pili (white bars), 1-2 pili (light gray bars), or at least 3 pili (dark gray bars). *n* = 300 cells analyzed from three independent biological replicates for all samples and data are displayed as the mean ± SD. All strains with ectopic P_BAD_ constructs were induced with 0.2% arabinose. Parent and Δ*pilT* data from **Fig. 1C** are included for comparison. (**B**) Representative images of piliated cells from strains in **A**. Phase images (top) show cell boundaries and fluorescence images (bottom) show AF488-mal labeled pili. Strains that retain native *pilT* are denoted “+” and strains with Δ*pilT* mutations are denoted “-”. Scale bar = 1 μm.

**Fig. S2.**
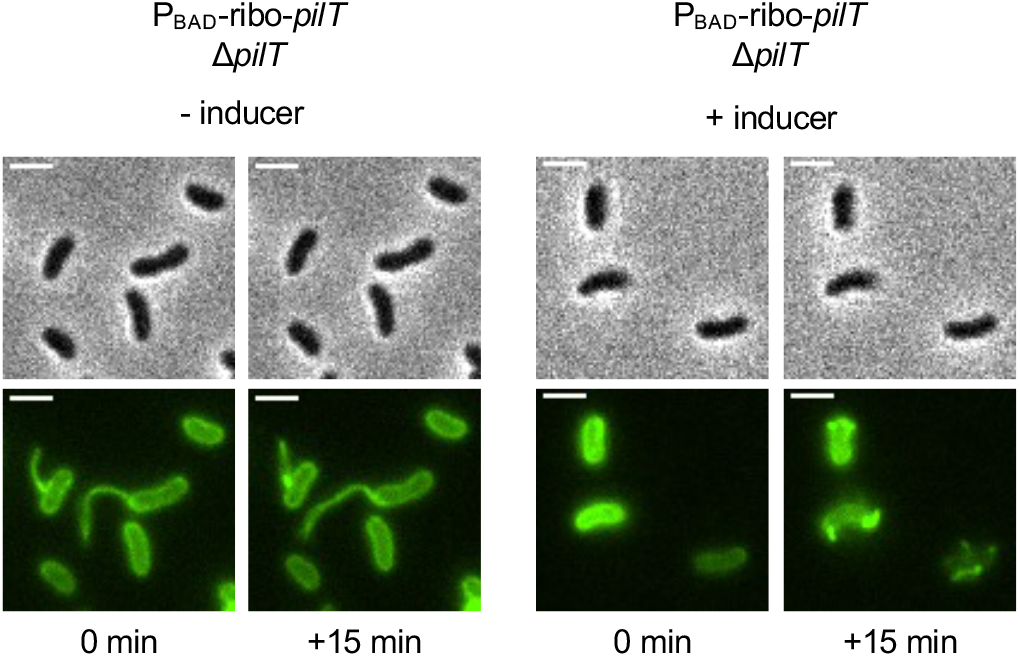
Induction of *pilT* rapidly recovers MSHA surface piliation. Representative images of a P_BAD_-ribo-*pilT ΔpilT* strain when *pilT* expression is rapidly induced under the microscope. Phase images (top) show cell boundaries and fluorescence images (bottom) show AF488-mal labeled pili. Samples were applied to a slide with a gelzan pad either lacking inducer (left; “-inducer”) or including 0.2% arabinose and 1.5mM theophylline as an inducer (“+ inducer”). Each sample was imaged immediately after application of the gelzan pad (“0 min”) and again after 15 min (“+15 min”). Scale bar = 2 μm.

**Fig. S3.**
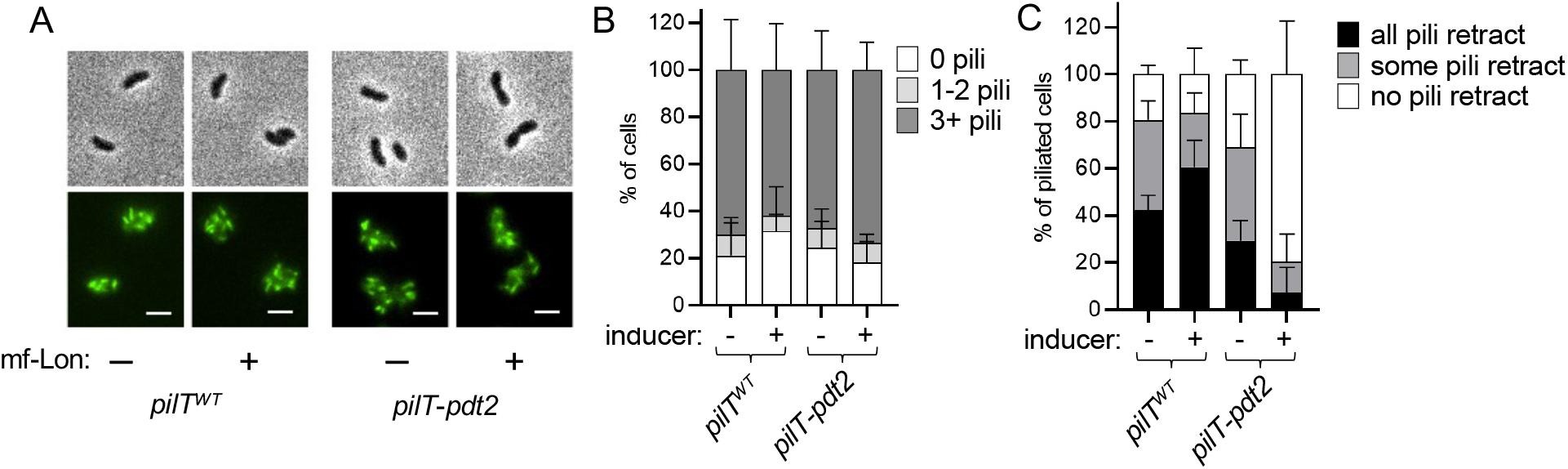
PilT is not required to maintain MSHA surface piliation. (**A**) Representative images of cells where mf-Lon was (“+”; 0.2% arabinose) or was not (“-”) induced. Phase images (top) show cell boundaries and fluorescence images (bottom) show AF488-mal labeled pili. Scale bar = 1 μm. (**B**) Quantification of piliation in samples from **A**. Cells were categorized as either having no pili (white bars), 1-2 pili (light gray bars), or at least 3 pili (dark gray bars). *n* = 300 cells analyzed from three independent biological replicates for all samples. (**C**) Quantification of imaging-induced retraction in samples from **A**. Graph displays the percentage of cells in each replicate that retracted all, some, or no pili during a three-minute timelapse. *n* ≥ 25 piliated cells analyzed for each of the three biological replicates. All data are displayed as the mean ± SD.

**Fig. S4.**
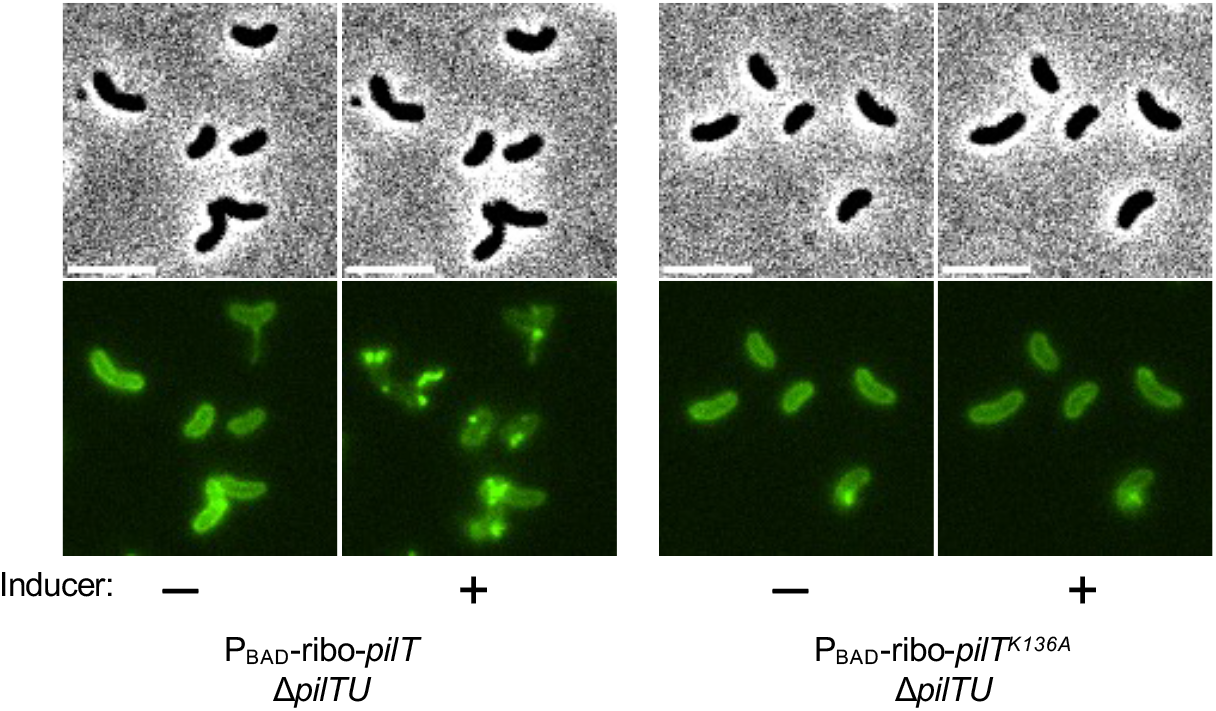
PilT does not require PilU to promote MSHA surface piliation as long as its ATPase activity is intact. Fluorescence microscopy of the indicated strains. Phase images (top) show cell boundaries and fluorescence images (bottom) show AF488-mal labeled pili. Each sample is shown before (ꟷ) and after (+) induction. Scale bar = 4 μm. Data are representative of three independent experiments.

**Fig. S5.**
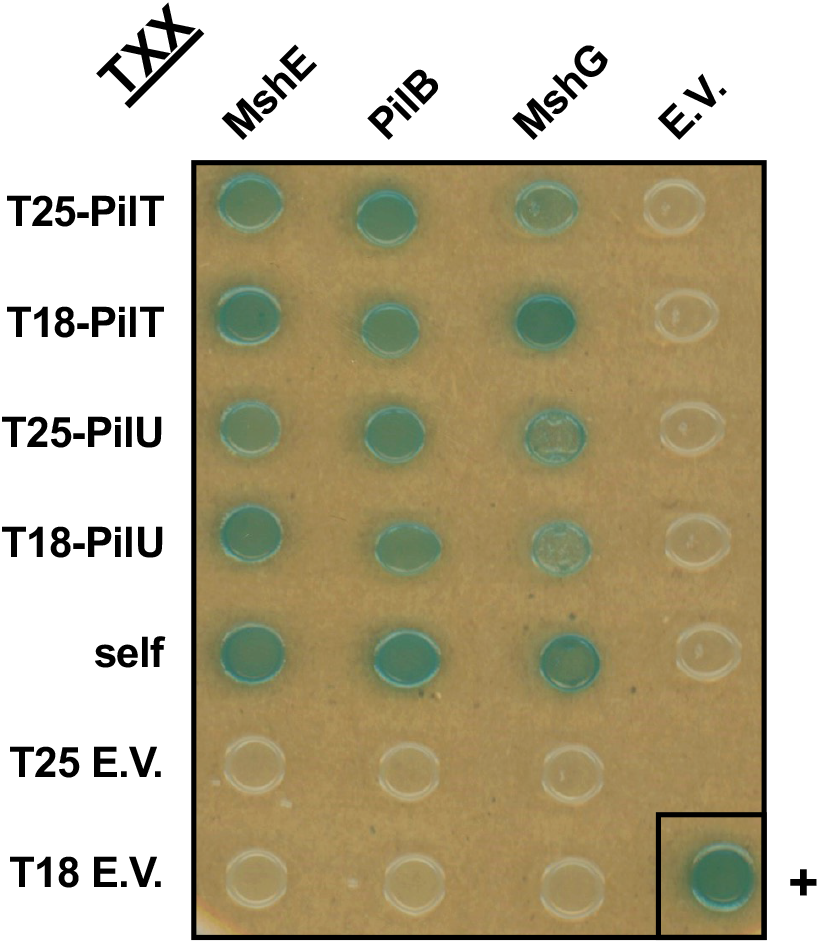
PilT and PilU directly interact with the MshE and PilB extension motors. Representative image of a BACTH assay between T25- and T18-fusions of the indicated proteins. Both PilT and PilU displayed strong interactions with themselves (“self”), the extension motors (MshE and PilB), as well as the MSHA platform protein (MshG). “E.V.” denotes a pairing with an empty vector, and “+” indicates the BACTH positive control (T25-Zip + T18-Zip). This image is representative of three independent experiments.

**Fig. S6.**
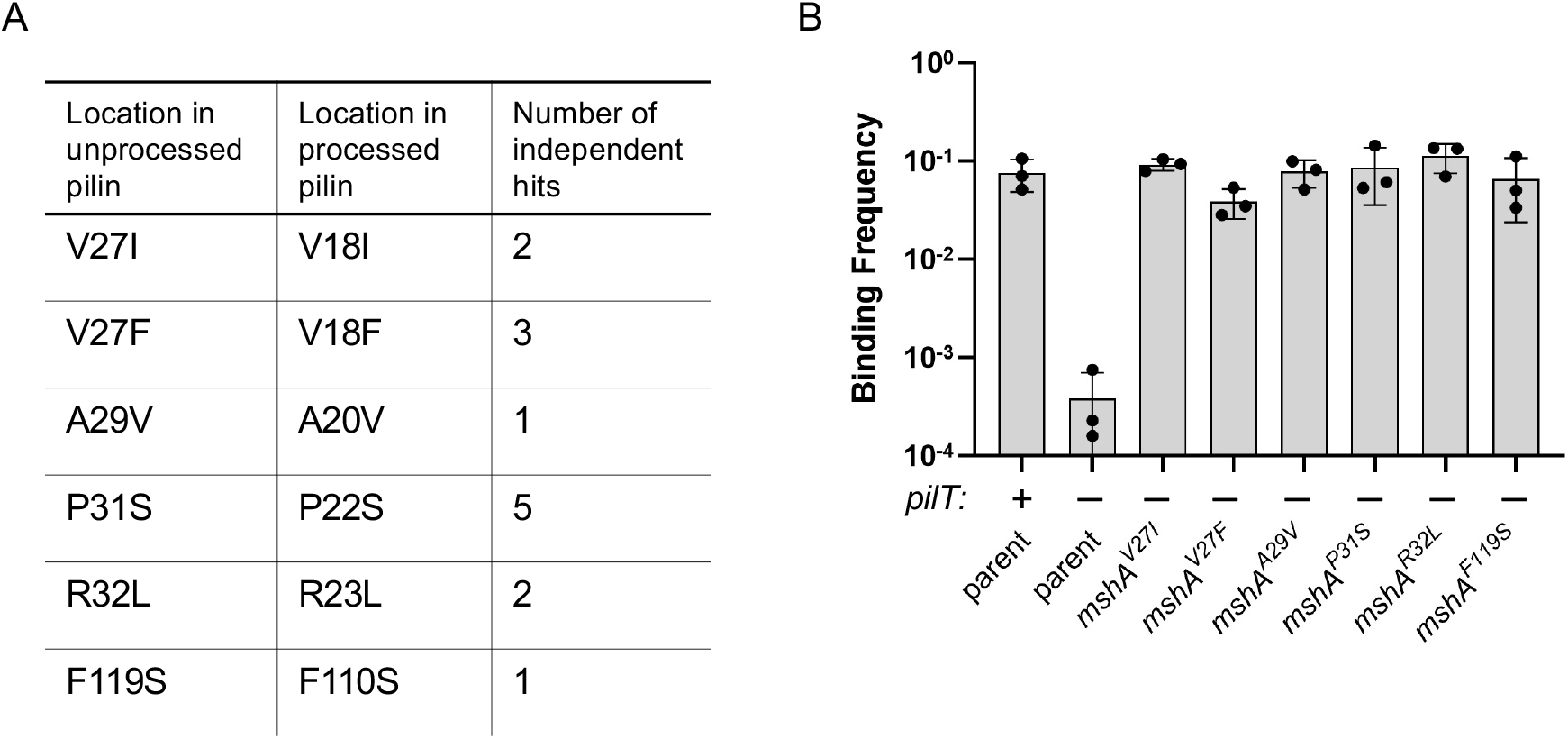
MshA suppressor mutants enhance adherence. (**A**) Table of the MshA mutations isolated in the Δ*pilT* suppressor screen. The relative position of the mutated residues is indicated relative to the MshA start codon (location in unprocessed pilin) and relative to the first amino acid in the pilin after being processed by the pre-pilin peptidase (location in processed pilin). “Number of independent hits” denotes the number of distinct genetic lines in which the indicated suppressor mutation was isolated. (**B**) Binding frequency of the indicated strains to the wall of culture tubes. Strains that retain native *pilT* are denoted “+” and strains with Δ*pilT* mutations are denoted “-”. Data are from three independent biological replicates and shown as the mean ± SD.

**Fig. S7.**
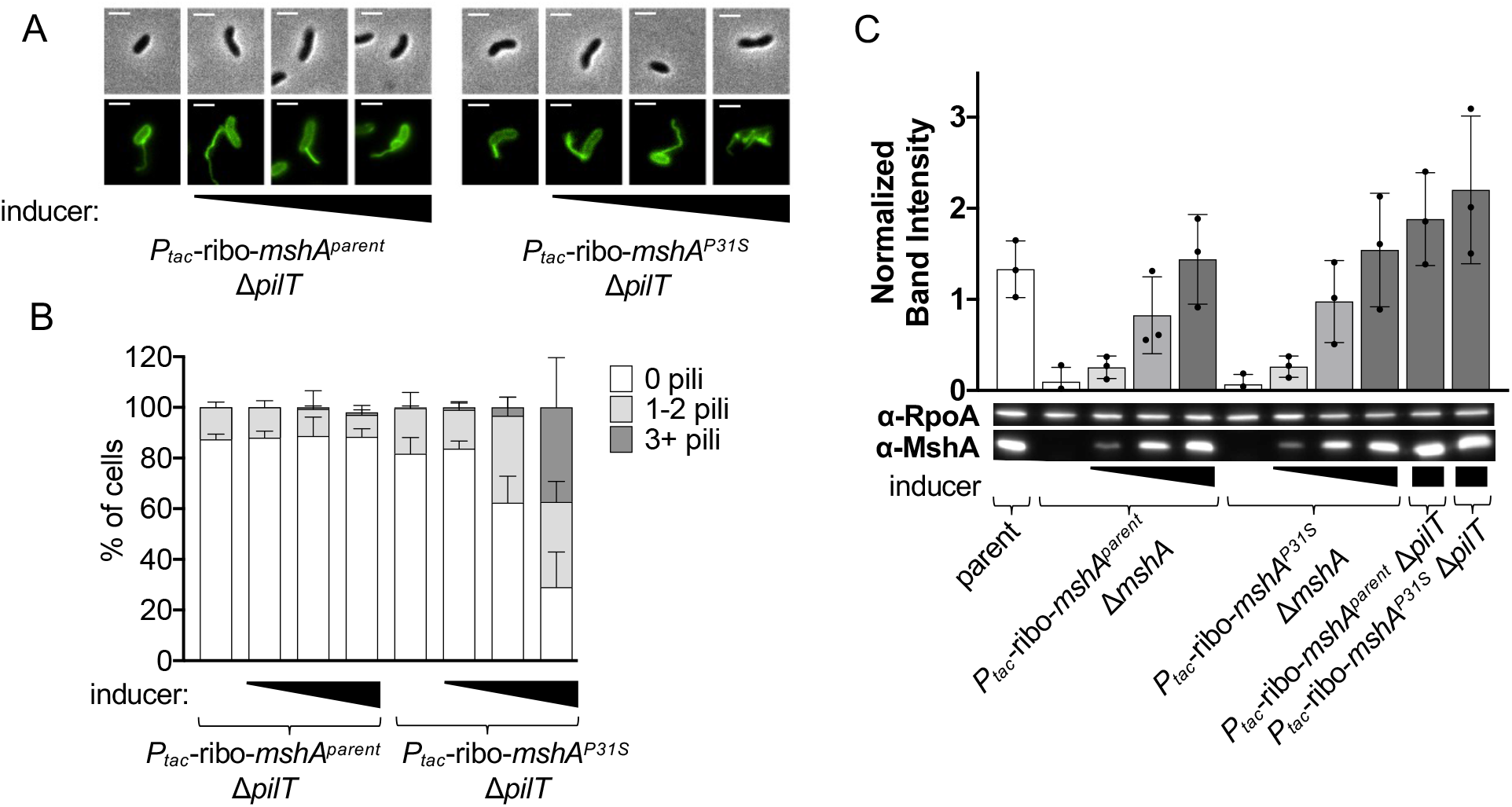
Full induction of MshA^P31S^ is required to recover piliation in the absence of *pilT*. (**A**) Representative images of piliated cells from strains under increasing induction of either P_tac_-riboswitch-*mshA^parent^* or P_tac_-riboswitch-*mshA^P31S^*. All *mshA* alleles in these strains (native and ectopic) contain the T70C mutation needed for AF488-mal labeling. Concentrations of inducer used from left to right: (1) 0 μM IPTG + 0 μM theophylline, (2) 5 μM IPTG + 75 μM theophylline, (3) 20 μM IPTG + 300 μM theophylline, and (4) 100 μM IPTG + 1.5 mM theophylline. Phase images (top) show cell boundaries and fluorescence images (bottom) show AF488-mal labeled pili. Scale bar = 2 μm. (**B**) Quantification of piliation in samples from **A**. Cells were categorized as either having no pili (white bars), 1-2 pili (light gray bars), or at least 3 pili (dark gray bars). *n* = 300 cells analyzed from three independent biological replicates for all samples. (**C**) Western blot quantification of cell-associated MshA in the indicated strains in the induction conditions used in **A** and **B**. Band intensities are normalized to the RpoA loading control. Data are from three independent biological replicates. All data are displayed as the mean ± SD.

**Fig. S8.**
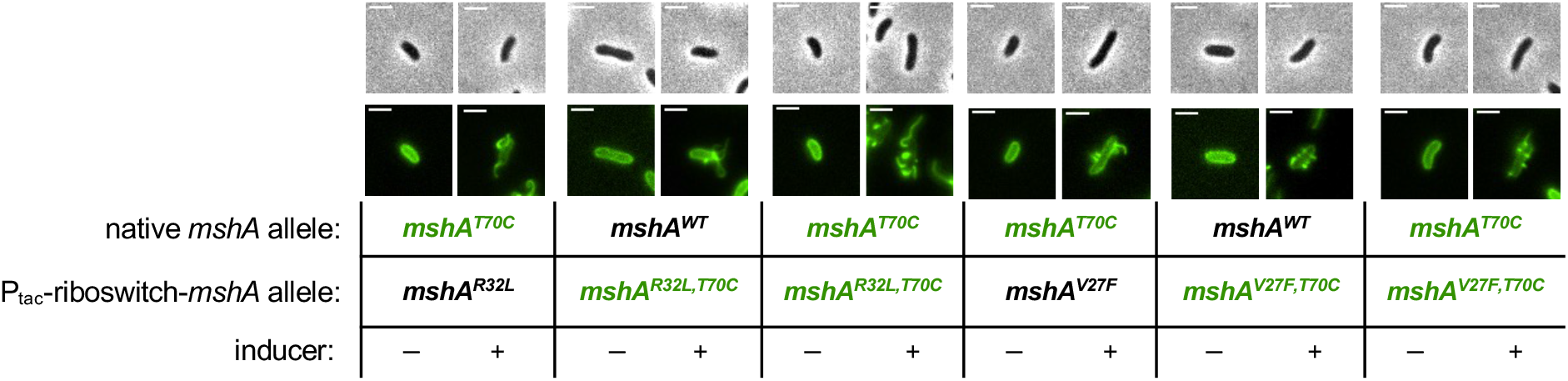
The MshA^V27F^ and MshA^R32L^ suppressor alleles promotes assembly of wild-type MshA. Representative images of cells from Δ*pilT* strains expressing the indicated *mshA* allele at the native locus (native *mshA* allele) and ectopic locus (P_tac_-riboswitch-*mshA* allele). Cells were either grown with (”+”) or without (”-”) 100 µM IPTG + 1.5 mM theophylline to induce the ectopic P_tac_-riboswitch-*mshA* allele in the strain as indicated. Only the alleles containing the T70C mutation can be labeled with AF488-mal and are denoted in green text in the table below the images. Scale bar = 2 μm.

**Table S1:**
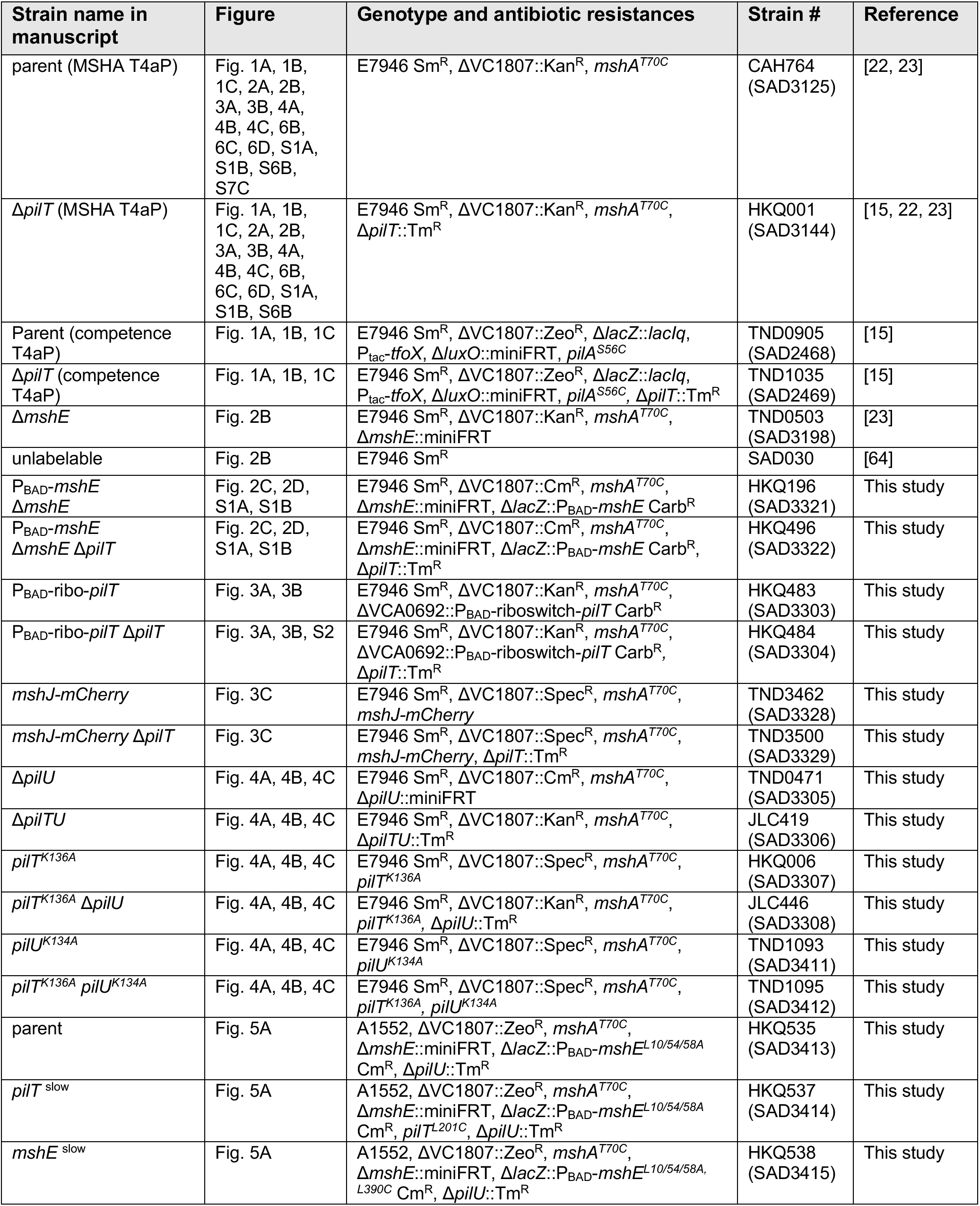
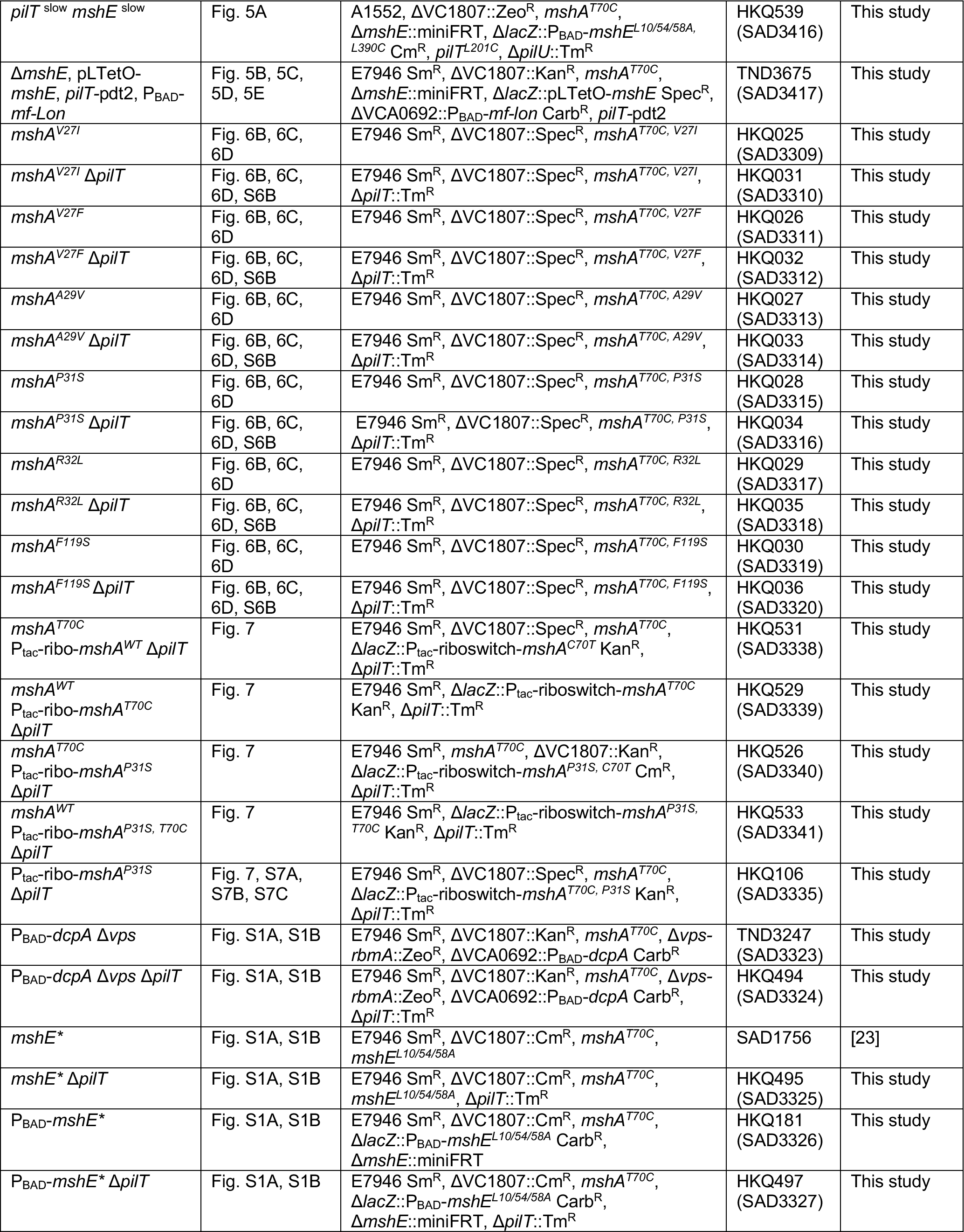
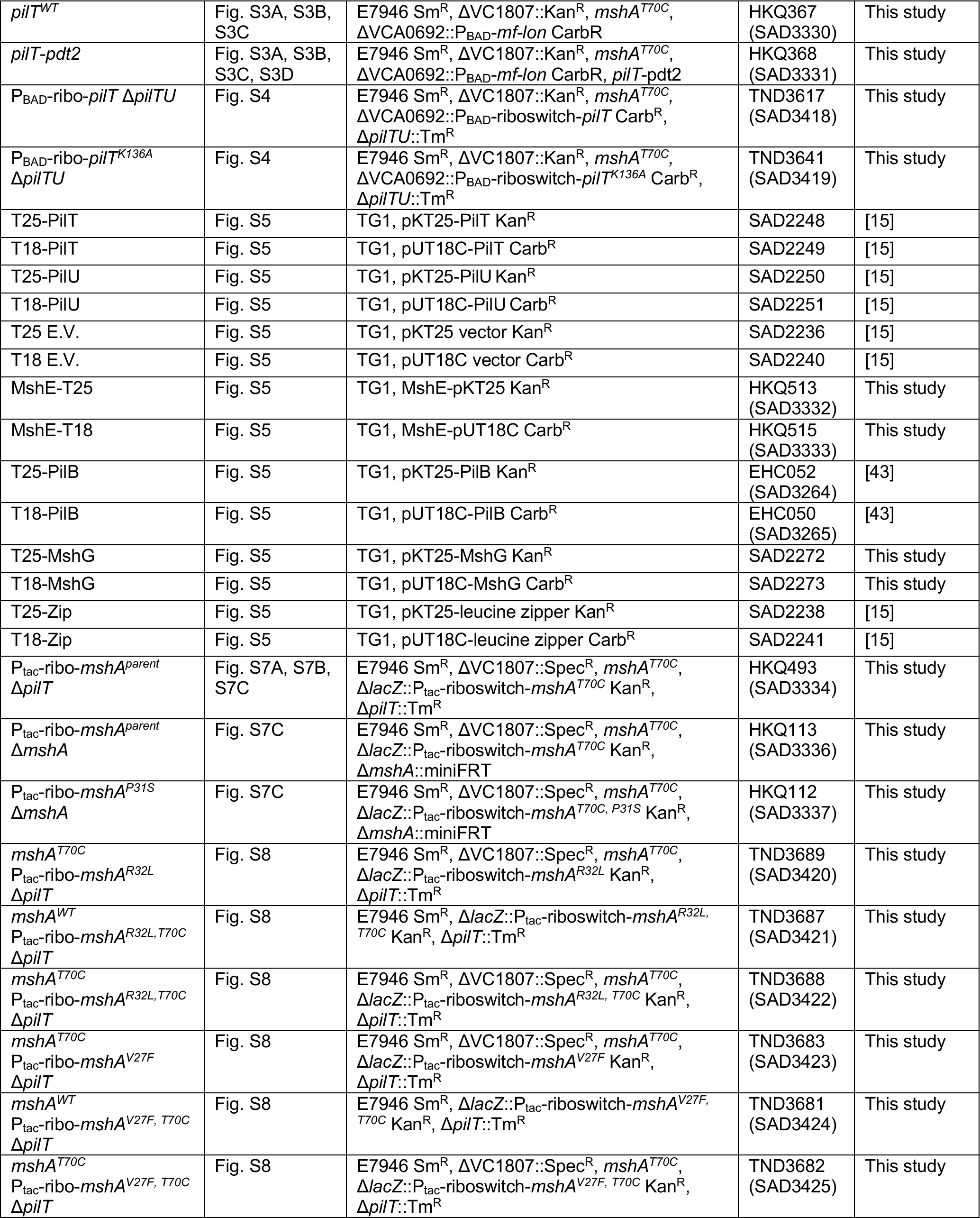
Strain List.

**Table S2:**
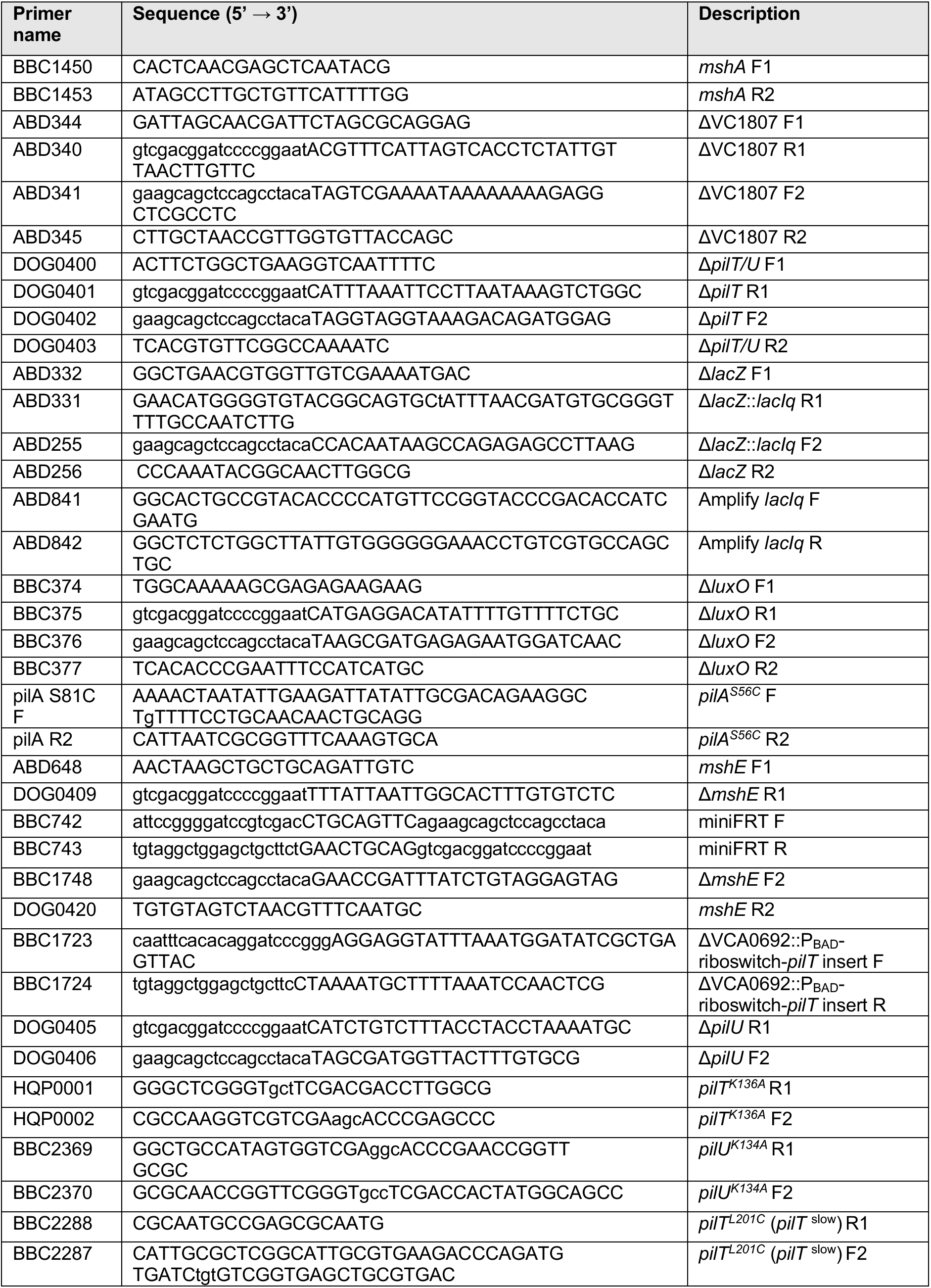
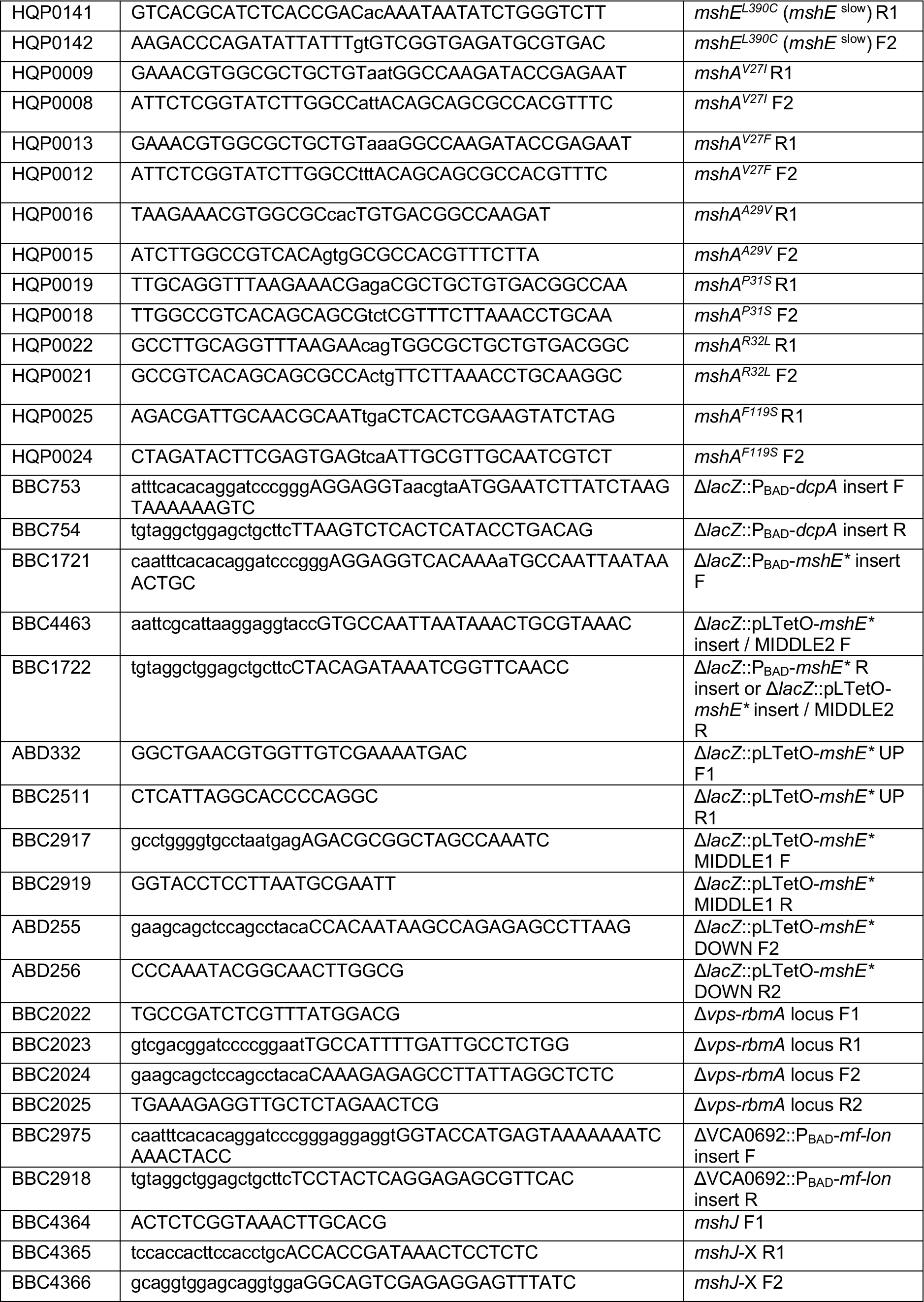
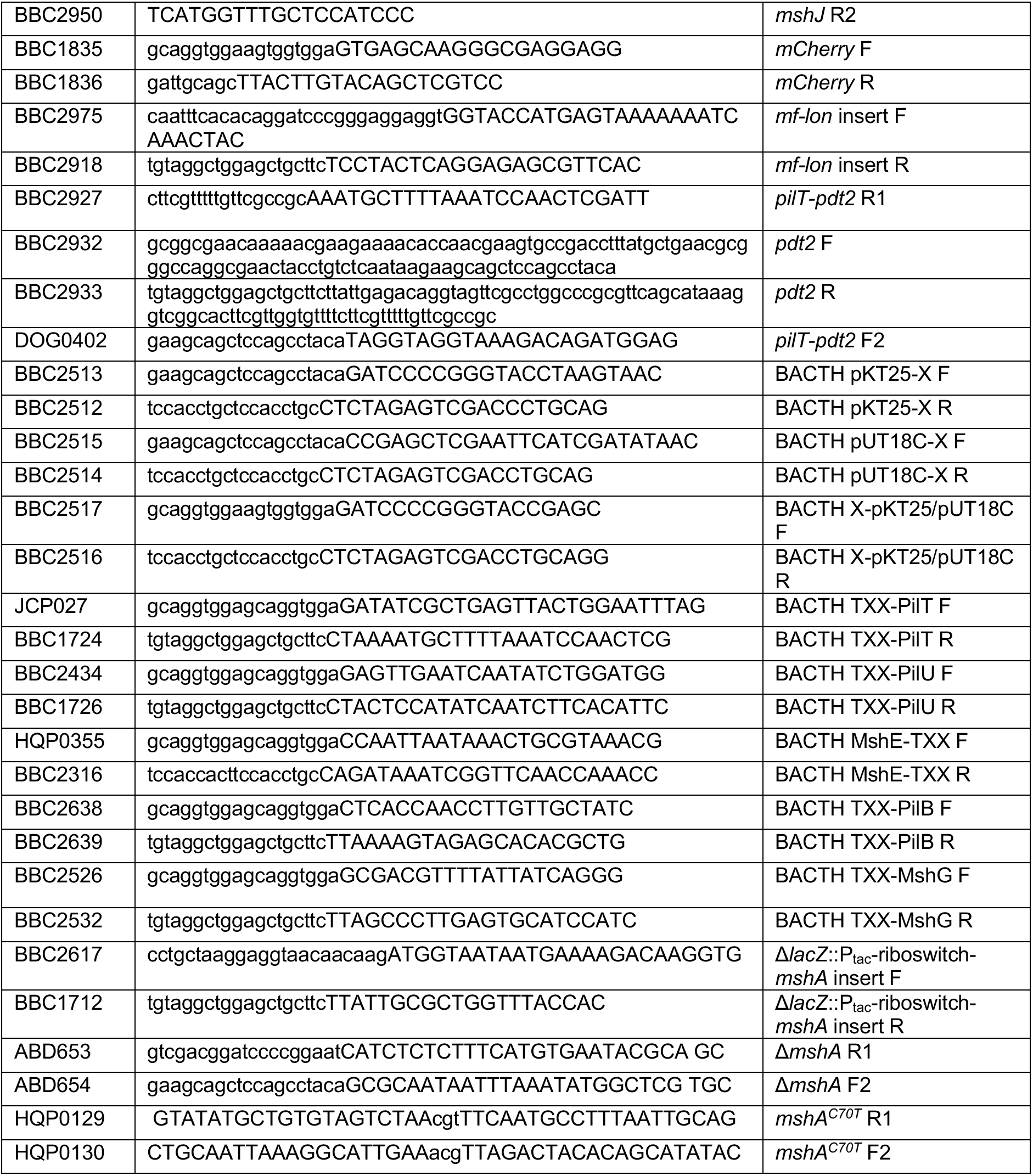
Primer List.

## Notes

### Competing Interest Statement

The authors have declared no competing interest.

### Summary of Updates

.

